# A novel high-accuracy genome assembly method utilizing a high-throughput workflow

**DOI:** 10.1101/2020.11.26.400507

**Authors:** Qingdong Zeng, Wenjin Cao, Liping Xing, Guowei Qin, Jianhui Wu, Michael F. Nagle, Qin Xiong, Jinhui Chen, Liming Yang, Prasad Bajaj, Annapurna Chitikineni, Yan Zhou, Yunxin Yu, Jiang Xu, Xiaojun Nie, Lin Huang, Shengjie Liu, Jan Šafář, Hana Šimková, Weining Song, Baozhu Guo, Shilin Chen, Jaroslav Doležel, Zhaodong Hao, Qiang Cheng, Jianguo Liang, Jiansong Tang, Aizhong Cao, Qiang Wang, Xiangqian Lu, Shouping Yang, Hongxiang Ma, Jiajie Liu, Xiaoting Wang, Hong Zhang, Zhonghua Wang, Wanquan Ji, Changfa Wang, Fengping Yuan, Jisen Shi, Rajeev K. Varshney, Zhensheng Kang, Dejun Han, Haibin Xu

## Abstract

Across domains of biological research using genome sequence data, high-quality reference genome sequences are essential for characterizing genetic variation and understanding the genetic basis of phenotypes. However, the construction of genome assemblies for various species is often hampered by complexities of genome organization, especially repetitive and complex sequences, leading to mis-assembly and missing regions. Here, we describe a high-throughput gold standard genome assembly workflow using a large-scale bacterial artificial chromosome (BAC) library with a refined two-step pooling strategy and the Lamp assembler algorithm. This strategy minimizes the laborious processes of physical map construction and clone-by-clone sequencing, enabling inexpensive sequencing of several thousand BAC clones. By applying this strategy with a minimum tiling path BAC clone library for the short arm of chromosome 2D (2DS) of bread wheat, 98% of BAC sequences, covering 92.7% of the 2DS chromosome, were assembled correctly for this species with a highly complex and repetitive genome. We also identified 48 large mis-assemblies in the reference wheat genome assembly (IWGSC RefSeq v1.0) and corrected these large mis-assemblies in addition to filling 92.2% of the gaps in RefSeq v1.0. Our 2DS assembly represents a new benchmark for the assembly of complex genomes with both high accuracy and efficiency.

## Introduction

High-quality reference genome sequences are critical resources for the characterization of genomic structure and function, as well as the heritable phenotypes driven by underlying genomic variation. These resources are being applied in diverse areas of research, including genetic analysis^1^, crop improvement^2^, medical screening^3^ and synthetic biology^4^. Therefore, the genomics research community is highly motivated in working towards the objective of developing high-quality genome sequences. In practice, due to the difficulties associated with obtaining a fully assembled, refined genome, it is feasible to achieve a nearly complete level of semi-contiguous assembly, with many genomic regions covered by a small number of contigs, but there remain gaps, mis-assemblies and masked regions of low sequence quality or with repetitive sequences^5^. Such sequences provide essential references that can be used for whole-genome comparison, evolutionary and phylogenetic analyses^6, 7^, identifying natural variants or key genes^8–11^, and genome-wide transcriptome analysis^12^. However, the reliance on a single reference genome can result in the inability to identify large structural variations in different genetic backgrounds, such as insertions and duplications; accordingly, recent studies have attempted to overcome this limitation by comparing *de novo* assemblies of each genotype—a collection termed a “pan-genome”^1, 13, 14^.

With the invention of long-read sequencing technologies such as single-molecule real-time (SMRT) sequencing^15^ and Nanopore sequencing^16^, the production of highly contiguous reference genomes for species with small genomes of relatively low complexity has become increasingly practical. However, there remain major challenges for species with large or complex genomes, such as those with repetitive sequences^17^ and high-order polyploidy^18^, including polyploids with a high degree of heterozygosity^19, 20^. Long repetitive sequences are found in the genomes of almost all eukaryotes, especially species with large or polyploid genomes such as gymnosperms^21^ and species of the Poaceae family^22^. The highly complex and repetitive nature of these genomes results from chromosomal structural variation due to unequal cross-over^23^, segment replication^24^ and insertion of various transposable elements^25^, among other modes of recombination and mutation during the course of evolution. Such repetitive regions often result in mis-assembly and reduce the quality of the reference genome, thus adversely affecting the accuracy of subsequent resequencing and pan-genomic analysis in these regions^26^. Another example is the dikaryon or polykaryon genome characteristics of some basidiomycetes, in which two or more cell nuclei with similar genomes can coexist in a single cell, leading to difficulties in distinguishing genome-wide repeats and resolving sequences of all chromosome sets within a single strain^27, 28^.

Mis-assembly is a common result of difficulties in assembling genome regions that are complex and repetitive. In a mis-assembly event, two or more contigs or scaffolds are erroneously joined when they in fact originate from non-adjacent positions, including positions on different chromosomes. For assemblies using short reads, mis-assembly often results from the discarding of contigs with short lengths. In contrast, for assemblies based on long reads alone or utilizing hybrid sequencing with both long and short reads, mis-assembly is more likely to result from errors or losses in overlap detection due to sequencing errors of long reads^15, 29^. During the construction of scaffolds, mis-assembly can originate from incorrect connections that stem from a small proportion of weak, errant or conflicting lines of evidence from major long-range technologies, such as mate-pair sequencing^30^, Hi-C^31^, optical mapping^32^, 10x genomics technology^33^, genetic maps, and bacterial artificial chromosome libraries (BAC libraries) or fosmid libraries, carrying a particularly high risk when connecting contigs with short lengths or incorrect structures. Following the initial assembly is often a stage of error correction, a highly complicated task in which corrections are made to as many mis-assemblies as practical, with direct costs and labour expenses far exceeding those of initial assembly^34^. Therefore, refinements in sequencing and assembly algorithms are essential prerequisites for reliable assembly of highly complex genomes^35^.

While next-generation shotgun sequencing methods are associated with a high risk of mis-assembly, this risk is reduced for “gold standard” assembly workflows using map-based approaches, including chromosome sorting^36^, BAC libraries, physical mapping^37^, and clone-by-clone sequencing, among others. These map-based approaches reduce the problem of genome assembly from the whole-genome scale to the scale of a BAC clone or another chromosomal fragment. Thus, regional reference sequences are produced and then used for either *de novo* assembly or error correction following whole-genome sequencing (WGS). By using this strategy, gold standard assemblies were successfully produced for several key species of major academic or economic value, including humans^38^, rice^39^, maize^40^, wheat^41^ and Arabidopsis^42^. However, these gold standard approaches tend to be highly laborious and are often cost prohibitive, which severely limits their usage. An impressive example is the wheat reference sequence (RefSeq) v1.0, the product of 13 years of collaborative interdisciplinary research coordinated by the International Wheat Genome Sequencing Consortium (IWGSC), involving collaboration among over 200 scientists from 73 research institutes in 20 countries^41^. RefSeq v1.0 consists of over 15 giga base pairs (Gb) with over 85% of the genome consisting of repetitive sequences. While this accomplishment established a new precedent for the robust assembly of complex genomes, its quality is limited to the extent that the assembly still contains 522,751 gaps and 481 mega base pairs (Mb) of unanchored contigs in defined chromosome assemblies.

Here, we describe a workflow with innovations in pooled sequencing and assembly algorithms that enable highly robust and accurate assembly of reference sequences with reduced labour and material costs. We designed a scalable two-step system for pooling samples and hybrid sequencing, allowing sequencing data for hundreds or thousands of BAC clones to be easily obtained in a single experiment. We developed the Lamp assembler to process pooled short reads along with pooled long reads and produce a complete assembly for BAC clone sequences. To demonstrate the capacity of our workflow for assembly of complex genomes, we chose the short arm of wheat chromosome 2D (2DS) as a benchmark. Our workflow reduces the sequencing cost to less than $10 per BAC clone and yields gapless assemblies for 98% of BACs. The contiguous 2DS contigs were produced with a contig N50 size of 1.1 Mb. Our workflow significantly improves the sequence accuracy of the 2DS portion of RefSeq v1.0, as we detected and corrected at least 48 large mis-assemblies and filled 92.2% of 10,434 sequence gaps. We validated the accuracy of our 2DS assembly by chromosome anchoring experiments using nullisomic-tetrasomic wheat lines^43^, as well as Sanger sequencing. These results were confirmed further by comparison of survey sequences specific to chromosomes or arms^44^, whole-genome profiling (WGP) data and physical maps^41^. Our Lamp algorithm exhibits significant potential to be incorporated into a wide range of low-cost sequencing solutions for genome projects of bacteria, monokaryon and dikaryon fungi and plant organelles, among other targets. In summary, our workflow is widely applicable across diverse species for the production of high-quality reference sequences.

## Results

### Scalable two-step sample pooling design and the Lamp assembler algorithm

We designed a two-step pooling system for high-throughput production and cross-referencing of short- and long-read data with multiplexed samples. Short reads were generated from the primary pools, which were further pooled to form “super pools” from which long reads were generated (Figure 1a, Materials and Methods). Primary pools from different types of samples are compatible within a super pool, thus providing throughput advantages and reducing per-sample sequencing costs while maximizing the flexibility of sample preparation for long-read sequencing. The pooling design reduced the sizes of sequencing libraries and did not require the sequencing barcodes that are commonly used to distinguish reads across primary pools or clones, which improved the platform-level flexibility of the pooling design and reduced the associated costs and labour.

**Figure 1.**
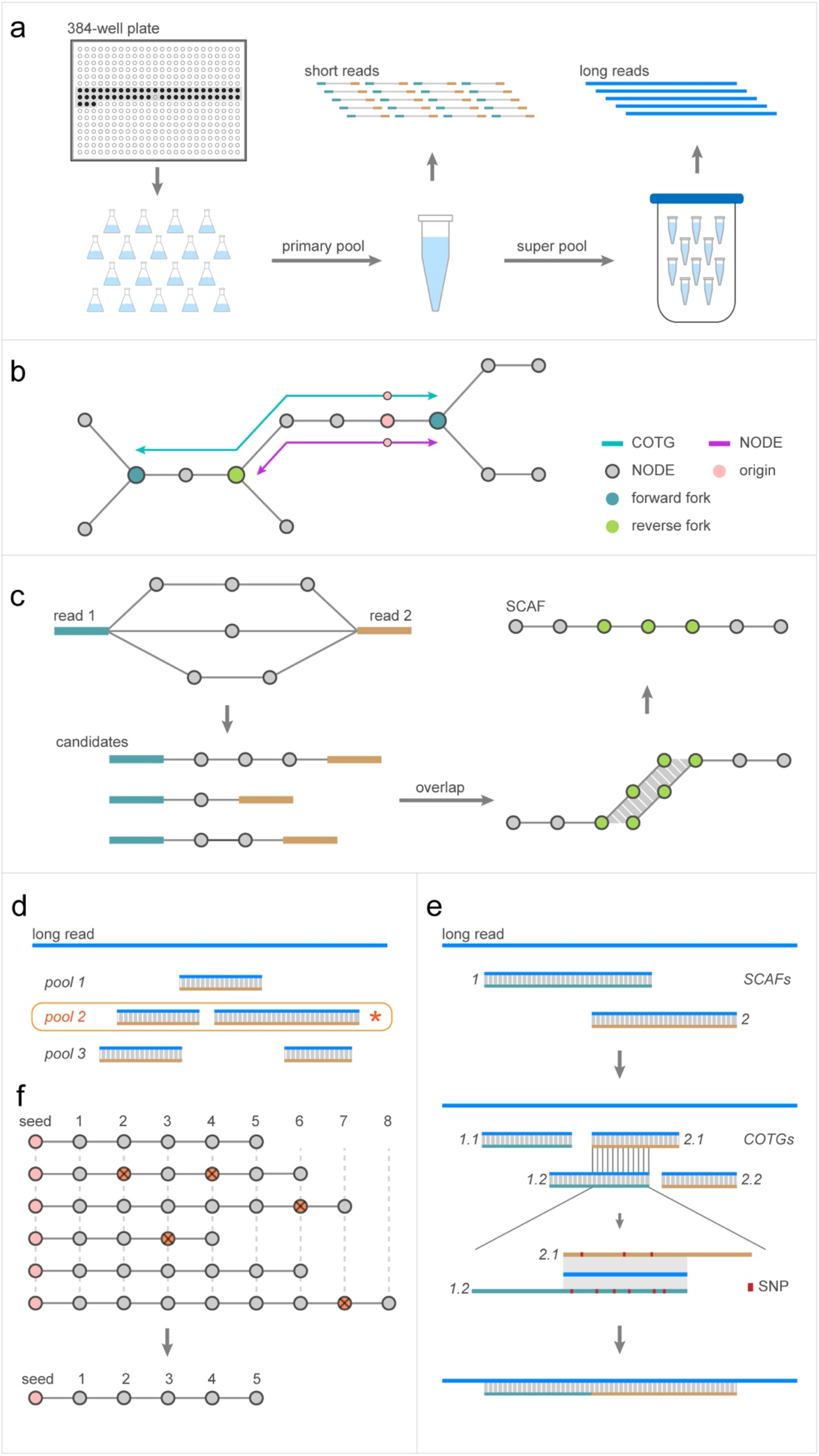
Overview of the pooling design and the Lamp assembler algorithm: **(a)** BAC clones were selected and cultured individually in Erlenmeyer flasks. The cultures were combined into mixtures with equal concentrations of cells from each culture, followed by plasmid extraction for short-read sequencing. These samples were further pooled into equal-concentration mixtures of combined plasmid DNA primary pools, producing super pools for long-read sequencing. **(b)** A given *k*-mer was extended by following a *de Bruijn* graph. The extension was suspended upon reaching a fork point to produce a unitig for the NODE contig set and continued until reaching a forward fork to produce a COTG contig. **(c)** Between reads of each read pair, paths were extended to fill the inner gap, and the outer ends of the reads were extended as COTG contigs. Overlaps were detected between contigs, and overlapping contigs were merged to produce SCAF contigs. **(d)** Alignments of a given long-read segment against SCAF contigs from multiple primary pools were compared to determine the source of long reads. A given long read was assigned to the primary pool with which alignment produced the greatest total aligned sequence length. **(e)** Alignments of long reads to SCAF contigs were first converted to alignments with COTG contigs and then compared with each other. The retained alignments were further downgraded to NODE-based chains connected by long reads. Inner gaps of these chains were filled first by attempting path extension and subsequently filled with the candidate extension deemed optimal based on the percent identity of alignments. **(f)** Seed contigs were used as scaffolds against which long-read chains were aligned, producing gapless matrices with each seed as an origin. Finally, these seeds were extended in accordance with these gapless matrices, yielding the final set of contigs on a chromosome scale.

Lamp is a *de novo* hybrid assembler algorithm designed to split long reads and assemble contigs using long reads as well as *k-*mers from short reads (Figure 1). The entire assembly process does not rely on reference genomes, physical maps, Hi-C data, optical maps, etc. It benefits from the dual advantages of long-read and short-read sequencing; high-accuracy short reads of sufficient coverage provide assurance of sequence integrity, while higher-order structural integrity is strengthened by long reads.

For each primary pool, Lamp constructs a *de Bruijn* graph^45^ from 99-mers disassembled from short reads and subsequently generates three contig sets of varying lengths and accuracies, termed NODE, COTG and SCAF, respectively (Figure 1b and 1c, Materials and Methods). In short, NODE is a set of unitigs of the highest confidence because they are produced by greedy extension—for any given extension, there is only a single candidate. The unitigs in NODE are 99 base pairs (bp) of minimum length, with a maximum of 98 bp overlap between unitigs. COTG is generated by extension of NODE. COTG construction begins with extension of each unitig along the *de Bruijn* graph, on both the 5’ and 3’ ends, until a dead end or a forward fork is reached. COTG has a structural accuracy comparable to that of NODE while featuring greater contig and overlap lengths. SCAF is produced by extension of gap-filled read pairs. For each given read pair, the *de Bruijn* graph was traversed to identify all candidate extension paths that could fill the inner gap. When such a candidate or candidates were found, one or more corresponding filled chains of unitigs were produced. Lamp next applies a step-by-step extension to fill chains on both ends, detecting and utilizing their overlaps to select candidates of the longest overlap length at each step, ultimately producing SCAF contigs. SCAF contigs feature sequences of greater average length than NODE and COTG and with reduced confidence relative to the other two contig sets.

To determine the primary pool from which each long read originates, Lamp compares alignments of a given long read against all SCAF contigs produced from the corresponding primary pools and makes a judgement based on the total length of retained alignments after removing false positives (Figure 1d, Materials and Methods). For a given primary pool, Lamp uses each long read as a reference axis and generates a connection chain of aligned unitigs. For each aligned unitig, Lamp records the order, orientation and distance from adjacent unitigs (Figure 1e). SCAF is the first contig set to be aligned to the long reads because the length superiority of SCAF provides an advantage for the rate of successful alignment of short unitigs to long reads. The initial alignments are next transformed to substitute SCAF for COTG, thus avoiding the influence of structural issues in SCAF. Lamp sequentially compares each alignment to a given portion of the long read and then removes alignments or portions deemed likely to be false positives based on identity percentages and the number of base mismatches. The initial unitig chain is obtained after the alignments are further converted to NODE-based alignments. Lamp again traverses the *de Bruijn* graph to find all extension path candidates that may fill the gaps in the chain and selects the optimal candidate based on comparison to long reads and generates long-read chains.

Lamp uses a greedy extension strategy^46^ to assemble genome sequences for each primary pool (Figure 1f). At the beginning of each extension loop, a non-repetitive unitig that is manually selected based on its length, coverage and forked interruption from a trial extension is used as the seed for extension. Repetitive unitigs are discarded when detected based on the frequent occurrence of conflicts during extension. Long-read chains in which a given seed is included are extracted, aligned using the seed as the origin, and extended to produce the genome chains of unitigs. When two or more extension path candidates appear, the extension pauses and then continues after manual judgement to select the most appropriate candidate.

Upon completion of extension, a terminal unitig or sub-chain is manually selected as a new seed to begin the next extension cycle. The loop is terminated upon reaching a telomere or BAC vector sequence or when the chain cannot be extended further. During BAC sequence assembly, Lamp checks for consistency between the sequence’s two terminal sub-chains and sub-chains at both ends of the vector in the long-read chains. Finally, gaps in the genome chain are filled using the self-corrected consensus sequence of long reads, producing the final genome sequence.

### Contiguous assembly and chromosome anchoring

For validation of our approach, we applied the two-step BAC pooling strategy to assemble contig sequences of the 2DS of the wheat cultivar Chinese Spring using the minimum tiling path (MTP) BAC library named TaaCsp2DSMTP. This MTP library comprises 3,025 BAC clones stored in eight 384-well plates and is a subset of the library TaaCsp2DShA (43,008 BAC clones with an estimated average insert size of 132 kilobases (kb)) used for WGP analysis (Figure s1a, Material s8)^47^. The 2DS WGP tags were previously used for the assembly of RefSeq v1.0^41^. In this work, a total of 60 primary pools were prepared for sequencing, each containing approximately 50 clones (Figure 1a, Table s1, Materials and Methods).

With the Lamp assembler, a total of 2,970 vector-to-vector BAC sequences were assembled to a gapless sequence (Table s2, Figure s1b, Materials s1, s2 and s3). The BAC sequences were 454.6 Mb in total length, with an average length of 153 kb and GC content of 46.5%, covering approximately 98% of the MTP clones (Table 1). Assembly could not be completed for some clones due to insufficient raw data resulting from *Escherichia coli* culture-related issues or due to structural complexity that exceeded the capabilities of the Lamp algorithm. We noticed that the lengths of wheat genomic sequences inserted in the vector differed among the primary pools (Table s1 and s3, Figure s2a), which is consistent with the source library being composed of two fractions with different insert sizes (Figure s2b, Material s8). We observed apparent plasmid replication errors mediated by the *E. coli* host. An example is the clone corresponding to the BAC_2_51 sequence, in which deletion of the 4,322-51,989 bp portion is supported by our long and short reads (Figure s3). The BAC sequences were further assembled to produce 458 contig sequences, with a total length of 271.4 Mb, average length of 593 kb and N50 length of 1.0 Mb (Table 1 and s4, Material s4). A total of 308 contigs were composed of two or more BAC sequences, while 150 contigs were derived from a single BAC sequence (Table s5). Three chimeric BACs were detected during contig assembly (Figure s4).

**Table 1.** Summary of assembly and chromosome anchoring

|  |  |
| --- | --- |
| Number of BAC sequences | 2,970 |
| Total length (bp) | 454,642,206 |
| Average length (bp) | 153,078 |
| GC content (%) | 46.5 |
| Number of contigs | 458 |
| Total length (bp) | 271,367,541 |
| Average length (bp) | 592,506 |
| N50 length (bp) | 1,039,371 |
| Number of 2DS contigs | 329 |
| Total length (bp) | 248,890,308 |
| Average length (bp) | 756,505 |
| N50 length (bp) | 111,7623 |
| Number of non-2DS contigs | 129 |
| Total length (bp) | 22,477,233 |
| Average length (bp) | 174,242 |
| Length of 2DS portion in IWGSC RefSeq v1.0 (bp) | 268,023,062 |
| Number of gaps | 10,434 |
| Number of mapped contigs | 326 |
| Total length of mapped contigs (bp) | 248,378,867 |
| Length of 2DS portions mapped by contigs (bp) | 248,373,015 |
| Coverage rate | 92.7% |
| Number of filled gaps in 2DS portion | 9,621 |
| Number of unfilled gaps | 813 |
| Closing rate | 92.2% |

The anchoring of contigs to chromosomes was determined by counting the lengths of each contig’s exact matches of at least 300 bp to the genomic survey sequences of all wheat chromosomes or arms (Table 1, s6 and s7). The results show that 329 out of the 458 assembled contigs with a total length of 248.9 Mb were anchored to the 2DS, while the N50 length increased to 1.1 Mb, and 290 contigs (88.1% of total contigs) originated from 2 or more BAC sequences. A set of 129 contigs with a total length of 22.5 Mb were scattered across genomic regions outside 2DS, including 111 (86.0%) assembled from one BAC sequence for each. These non-2DS contaminants were found in 169 BAC sequences, accounting for 5.7% of the assembled BACs. The locations of two contigs were particularly unclear. The first was the G_52_2 contig (128 kb), suspected to be located on the 2DS but with a similar match to the 2BL arm. The other was the G_519 contig (167 kb), located on a non-2DS contig with sequences matching both the 1BL and 5BL arms.

The chromosomal locations of non-2D contigs were verified by anchoring experiments using the nullisomic-tetrasomic lines of Chinese Spring wheat^43, 48^. Of 129 non-2DS contigs, 78 were anchored to the predicted chromosomes other than 2DS (Table s24, Figure s10). Failure to anchor for the remainder of the contigs may be attributable to weak specificity of primers (Figure s11). The anchoring results were further confirmed by comparison to RefSeq v1.0, and simultaneously, the exact positions of contigs in the chromosome assembly were obtained (Table s8). Out of 329 2DS contigs, 326 could be accurately anchored to the 2D chromosome assembly, while the locations of the T48, G_52_2 and G_528 contigs were undetermined. Two BAC sequences were found to be chimeric in this step, located at the ends of the G_185 and G_362_2 contigs, respectively (Table s5). At the chromosome level, the intervals covered by these contigs were distributed in the 0-268.0 Mb region of the 2D chromosome assembly, overlapping with the centromere region (264.4-272.5 Mb) determined by CENH3 ChIP-seq analysis^41^. In total, 92.7% of the 2DS assembly was covered by these assembled contigs (Table 1). This proportion is comparable to that in the 7DL physical map (92%) and greater than that in the 3B physical map (82%)^49, 50^. In addition, the contigs scattered across other genomic regions were all anchored accurately to the appropriate chromosome assemblies.

### Comparison of Lamp assembly results with the WGP tags and the physical map

Since each primary pool was pooled from known MTP clones, we assessed the correspondence between publicly available MTP clones and our assembled BAC sequences by matching these clones’ WGP tags to our BAC sequences according to their well positions in the plates received from CNRGV (Table s9). We first compared the MTP clones and our BAC sequences that originated from the same plate wells. The WGP tags were matched to 1,513, 1,011, and 446 BAC sequences with 100%, 99%∼80%, and less than 80% tags matched to MTP clones, respectively (Figure s5). From the sequences with less than 80% MTP tag matching, we found that eight pairs (15 BACs in total) could be matched to the same MTP clone for each member of the pair (Table s10). As an example, for the 46 WGP tags of the MTP clone DS.H059.M09 located in well I1 of plate No. 5, 19 and the remaining 27 tags were matched to BAC_37_34 and BAC_37_40, respectively. This is highly likely due to the original plate well containing the MTP clone having two different clones without overlaps. Taken together, the results show that corresponding clones were identified for 2,539 BAC sequences (85.5% of BAC sequences) (Table s9, Materials and Methods). In particular, among the five chimeric BAC sequences discovered during assembly as previously mentioned, all the sequences matched a unique clone for which all WGP tags could be retrieved, thus indicating that the chimeric status was not a result of mis-assembly during our workflow.

Considering that our BAC samples were pooled from the exact same plates of MTP clones from CNRGV (Materials and Methods), the low matching rates for the remaining 431 BAC sequences (terms unmatched BAC sequences) can presumably be attributed to cross-contamination in the MTP clone plates during clone selection. Notably, 413 out of 431 unmatched BAC sequences were concentrated among 24 primary pools: Nos. 17-24 and 41-56, corresponding to plate Nos. 3, 6 and 7 (Table s1 and s9). Accordingly, 422 of the 436 unmatched BAC clones on plate Nos. 3, 6 and 7 were selected from plate Nos. 25-32 and 81-88 of the source library (with a total of 112 plates) (Figure s1 and s6, Material s8). A total of 305 unmatched BAC sequences were re-anchored to the physical map with the assistance of adjacent BAC sequences in contigs (Figure s7a). Among these sequences, only 11 (3.6%) appear to overlap in position with BAC clones of the same primary pool for each primary pool in the physical map (Table s11, Figure s7b), further indicating the unmatched status of the remaining BAC sequences. When we tried to expand the matching range to all 3,025 MTP clones, 294 (68.2%) BAC sequences remained for which no match was identified using an 80% matching rate with tags as the threshold (Table s12 and s13, Figure s8), suggesting that the problem of unmatched sequences was not attributable to primary pool design and handling. As we attempted to expand the range to all 37,635 BAC clones with WGP tags in the source library, matching candidates were detected for all but 16 (3.7%) BAC sequences (Table s12 and s14, Figure s8), confirming that the problem was not due to errors caused by the Lamp algorithm. Moreover, the matching rates for plate Nos. 1, 2, 4, 5 and 8 were not unexpectedly low. The above results further support the robustness of our BAC sequence assembly.

Structural conformity to the physical map was evaluated using 199 contig sequences, each containing at least 5 MTP clones corresponding to unique BAC sequences (Table s15 and s16). A total of 183 contig sequences could be anchored to particular continuous physical map regions, providing validation for assembly in these regions. Each of the remaining 16 contig sequences corresponded to at least two physical map regions, a result mainly attributable to the Lamp algorithm’s accurate detection of missing or erroneous overlaps in the physical map (Figure 2a). For example, the T2 contig sequence, with a length of 2,089 kb, matched the ctg63 and ctg27 contigs in the physical map at the 1-1,051 kb and 1,043-2,089 kb regions, respectively. The sequences BAC_9_48 and BAC_5_12, corresponding to clones DS.H014.P24 and DS.H006.L03, were found to overlap with a length of 7.6 kb at the junction position (1,043-1,051 kb), although this overlap is not represented in the physical map.

**Figure 2.**
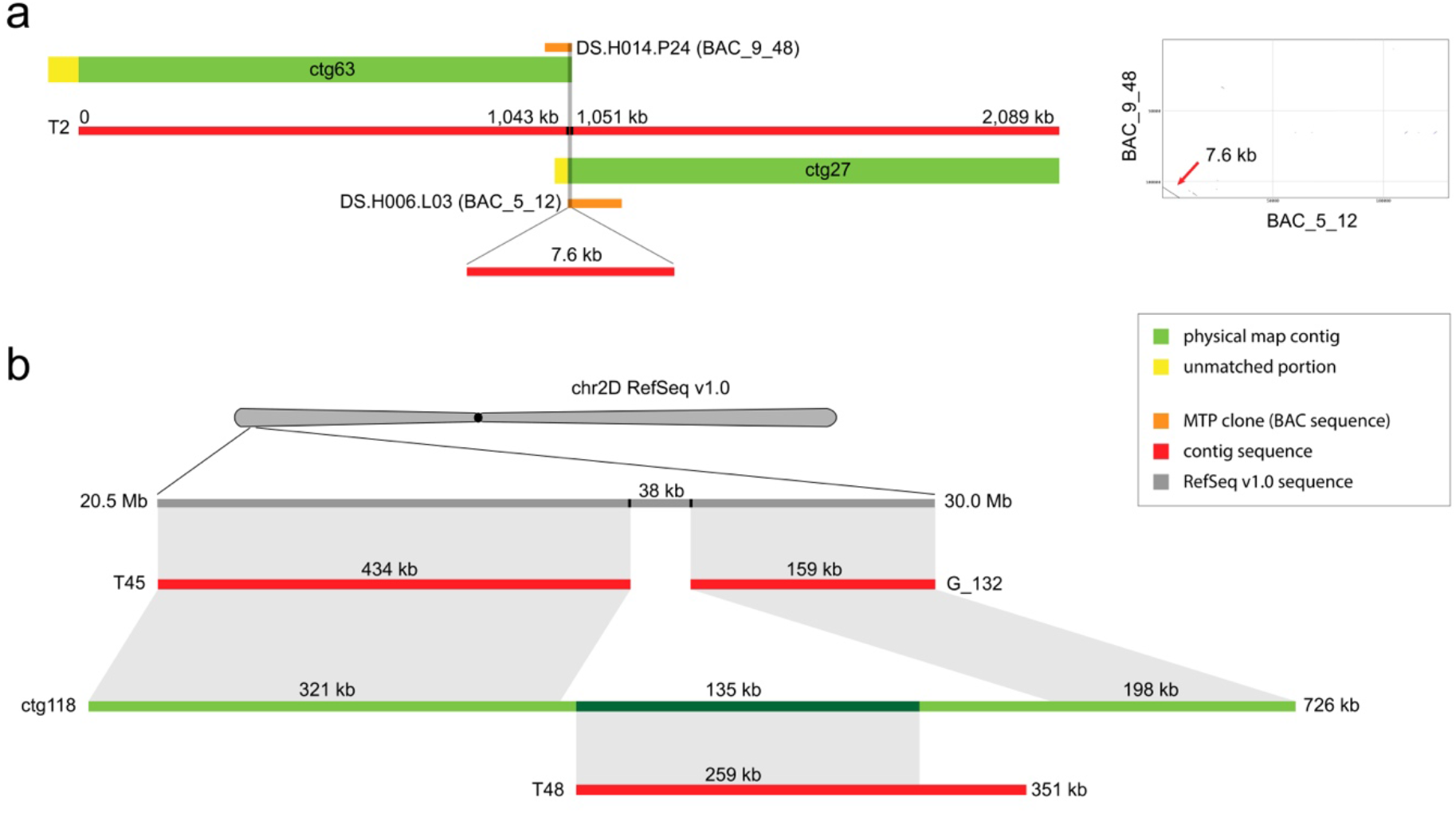
Comparison of contig sequences to physical map contigs: **(a)** Illustration of contig retrieval and dot-plot visualization showing the overlap that is absent from the physical map contigs but indicated by the assembly of the T2 contig sequence. **(b)** Illustration of a candidate anchoring position for the T48 contig sequence in IWGSC RefSeq v1.0 compared to physical map contigs.

Because the clones used to construct the physical map were not used to build RefSeq v1.0^41^, we then compared the consistency between the physical map and the RefSeq v1.0 assembly by making use of all contig sequences that overlapped at any given region in the continuous physical map (327 contig sequences in total). The results generally show agreement between sequences (Table s17). Notably, the unanchored T48 contig sequence (351 kb) was located in the ctg118 physical map contig between the T45 and G_132 contig sequences; thus, T48 was preliminarily anchored in the 20,795-20,833 kb (38 kb) region in the 2D chromosome assembly (Figure 2b, Material s6), suggesting potential for the Lamp assembler to resolve large mis-assemblies. Furthermore, a conflict was detected in the ctg71 physical map contig, in which two contig sequences mapped to chromosome 1B, but their distance from one another in RefSeq v1.0 reached 109 Mb.

### Revisions of large mis-assemblies and gaps in the IWGSC RefSeq v1.0

Our comparison revealed overall structural consistency between RefSeq v1.0 and our assembled contig sequences, further proving the reliability of our workflow (Figure 3). Despite this trend of general consistency, we detected 43 large structural inconsistencies on the 2DS assembly with interval lengths exceeding 20 kb, indicating the presence of large mis-assemblies in RefSeq v1.0 (Table s18 and Figure s9). This included 38 insertions, two deletions, and three inversions relative to RefSeq v1.0. These insertions involved contig portions of 3.9 Mb in total, with the largest interval at 602 kb (Figure 3). Deletions were found for two intervals of the RefSeq, with lengths of 506 kb and 82 kb. Among the three inversions, var_13 and var_36 were near the telomere, and var_21 was near the centromere. We also detected seven large structural differences on non-2DS contigs, including three insertions, one deletion, one local translocation and two large translocations relative to RefSeq v1.0 (Table s18 and Figure s9). We speculate that the two large translocations may be false positives resulting from chimeric clones, although this possibility remains uninvestigated. We attempted to predict gene models in these insertions (Table s19 and Material s5); a total of 292 putative genes were annotated, including *evm.model.T20.9,* which may encode a disease resistance-related protein, indicating that chromosomes assembled in RefSeq v1.0 may exclude some genes affecting key agronomic traits.

**Figure 3.**
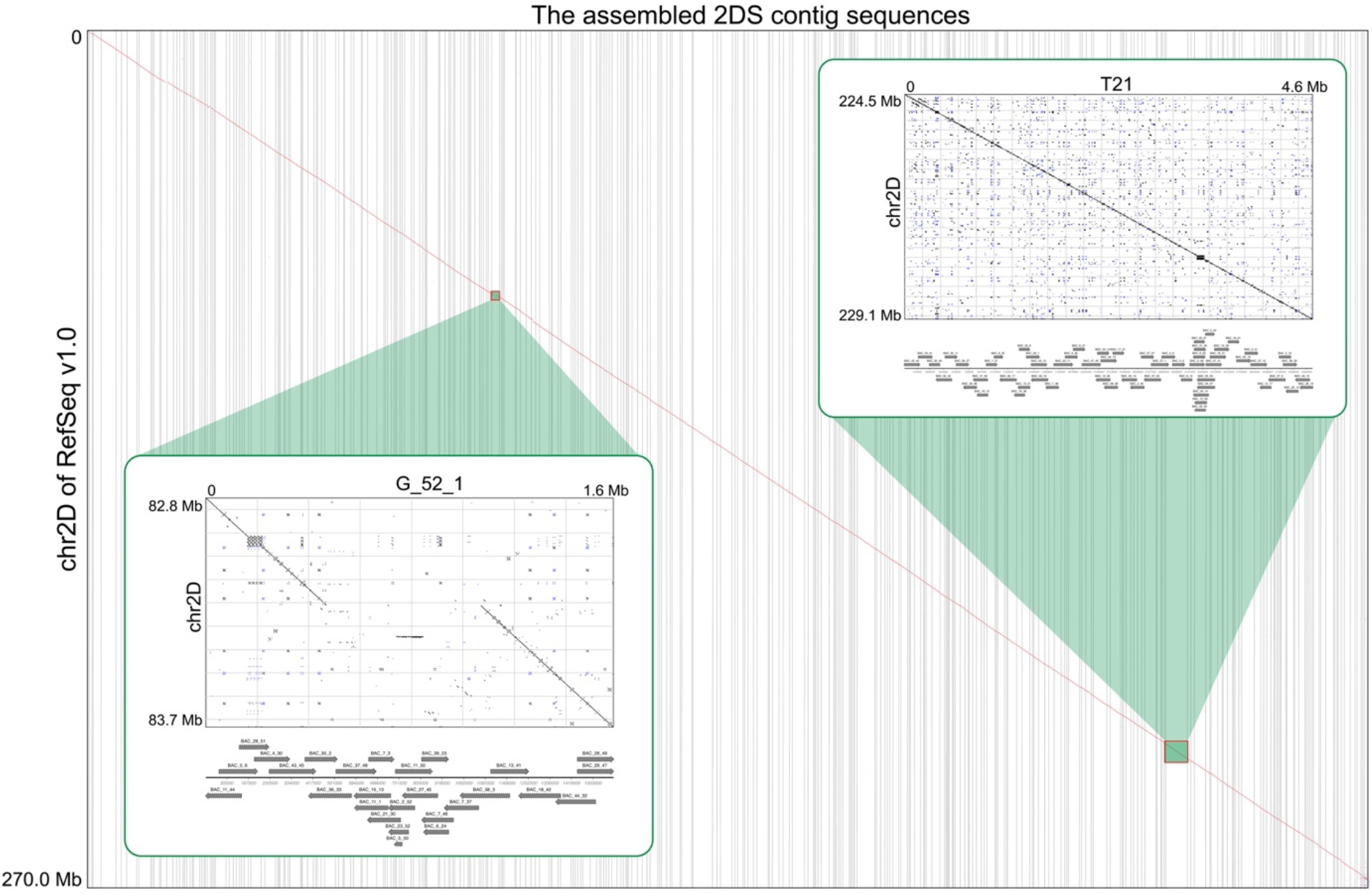
Global comparisons of our assembled 2DS contig sequences to IWGSC RefSeq v1.0. Dot-plot visualizations and assembly details of T21 and G_52_1 contig sequences were magnified to be inspected. The T21 contig sequence shows a high degree of structural consistency with the associated fragment of 2DS, while a 602 kb structural inconsistency in the G_52_1 contig sequence is revealed.

Two independent experimental validations were conducted for these 2DS insertions. Twenty-seven allele-specific markers were designed using sequences from 38 2DS insertions. Thirteen of these 27 markers were successfully mapped to 2DS using the nullisomic-tetrasomic wheat lines as previously described (Table s24, Figure s10 and s11). To investigate the sequence and structural accuracy of mis-assembly corrections, we verified 39 boundaries of large mis-assemblies in 2DS by PCR amplification coupled with Sanger sequencing (Table s25, Figure s12 and s13). These two approaches together validated 29 out of the 38 2DS insertions. Using matching thresholds of 3 kb and 99% similarity, matching portions were detected for 34 out of all 41 insertions in the unanchored sequences (chrUN) of the RefSeq (Table s20 and s21). This finding suggests that inaccurate anchoring of some contigs led to exclusion of these contigs from chromosomes in RefSeq v1.0. The reliability of our workflow was further indicated by investigation of sequences exactly matched by insertions in any of the chromosome survey sequences, with chromosome anchoring of all insertions confirmed, with two exceptions, namely var_33 (22 kb) and var_35 (22 kb) (Table s22 and s23). Notably, many survey sequences that exactly match 2DS insertions appear in the chromosome arms 1BS, 2BL and 6AS, with frequencies of occurrence markedly higher than those for other chromosomes or arms. In particular, chromosome survey sequences exactly matching var_28 can be found in all chromosomes or arms. Considered together, these multiple lines of evidence suggest that long repetitive regions featuring similar sequences in different chromosomes were the primary cause of inaccurate anchoring during the RefSeq v1.0 assembly process.

Of the 10,434 inner gaps represented by ‘N’ in 2DS of RefSeq v1.0, 9,621 gaps were covered by our contig sequences and could be filled after integration of the RefSeq with our results, while only 813 gaps remained unfilled (Table 1). The 92.2% filling rate is comparable to the 92.7% coverage rate of contigs on 2DS, suggesting that the inability to fill the remaining gaps is mainly a result of incomplete coverage of relevant genome portions by the MTP library. Therefore, WGP tags may be used for further screening and sequencing of BAC clones spanning the uncovered regions, likely reducing the number of gaps in the 2DS assembly to below 100. As a by-product of our 2DS assembly, 827 gaps of non-2DS regions were covered by assembled non-2DS contigs and could also be filled during integration.

## Discussion

In this paper, we report the development of a high-throughput pooling methodology and the Lamp assembler algorithm and demonstrate the combined use of these methods for a gold standard assembly workflow applied to the 2DS of wheat. These advances contribute towards the goal of producing reference sequences for large and/or complex genomes at low cost, conserving labour costs for sample preparation and sequencing by taking advantage of pooling and hybrid sequencing. The average cost per BAC clone was below $10 for the 2DS MTP library used as a benchmark; furthermore, this cost can be reduced to less than $5 by making use of newer sequencing platforms. Ongoing reductions in sequencing costs will help to overcome cost as a limitation for widespread use of our workflow, in turn mitigating a major obstacle in producing gapless assemblies of very high coverage (>99%) from BAC clones. Therefore, our workflow will become increasingly accessible and valuable as gold standard genome workflows are used for an increasing number of accessions, with increasing demand for high accuracy and cost-efficiency^51^.

The tedious and time-consuming process of constructing physical maps is altogether avoided in our workflow since knowledge of BAC overlaps is not needed for pooling design or sequence assembly^52^. Even in the absence of physical map construction, the Lamp assembler is able to produce vector-to-vector gapless assemblies for 98% or more of BAC clones by enabling contig assembly using overlaps of BAC sequences alone. Our benchmark assembly using 2DS of wheat demonstrates that these advantages apply even to complex genomes. In our benchmark assembly, 2,970 BAC sequences from the wheat 2DS MTP library were assembled into 458 chromosome-scale contigs. The pooling design developed for this workflow offers flexibility as samples may be added or excluded in response to results from sequencing; for example, a primary pool that fails to yield reads during sequencing of a corresponding super pool (due to reagent failure or other issues) may later be sequenced as part of a subsequent super pool. In practice, a highly practical approach to building primary pools is to group clones from adjacent wells on a given library plate. Clones from different samples can be combined and subsequently divided following assembly. For challenging steps of assembly, the Lamp algorithm can reduce the occurrence of mis-assembly by allowing real-time manual judgement. Compared to popular workflows, including those with labour-intensive correction processes as well as black-box style full-automatic assembly tools such as Canu, FALCON, MaSuRCA, DeNovoMagic and TRITEX^41, 53–56^, our workflow demands the lowest total costs for the processes of sequencing, assembly and error correction (Material s7). Upon completion of contig assembly, the final reference sequence can be easily generated by integrating data from long-range technologies such as Hi-C, optical mapping and genetic mapping.

We evaluated the accuracy of our workflow using a benchmark of 2DS of wheat, which has proven highly difficult to assemble at reference quality using WGS data alone^41, 57–59^. As wheat is a key staple crop of global importance, improvements to the reference sequence may lead to major agronomic benefits by advancing the characterization and improvement of agronomically important traits such as yield and pest resistance. Our workflow enables the complete correction of at least 48 large mis-assemblies in IWGSC RefSeq v1.0. Notably, in accordance with a pre-publication data sharing agreement following the Toronto International Data Release Workshop standard, only five of these are revised completely in RefSeq v2.0 (Material s6). Although the assembly coverage was limited to 92.7%, constrained mainly by limited coverage of the MTP library, the number of gaps in 2DS RefSeq v1.0 was nevertheless reduced from 10,434 to 813. Corrections of these large mis-assemblies and gaps may improve the continuity and structural accuracy of 2DS assembly to such an extent that 2DS becomes the portion of IWGSC RefSeq closest to a gapless and complete assembly. Our workflow can be applied to any available chromosome-specific BAC and/or MTP library to fill gaps and correct mis-assembly throughout IWGSC RefSeq. Moreover, the detection of inconsistencies between WGP tags and the corresponding clones in the MTP library, as well as of errant or missing connections in the physical map, can help prevent introduction of these artefacts into subsequent studies. Overall, our workflow provides a low-cost, gold standard reference-quality assembly solution that can be applied feasibly not only to improve IWGSC RefSeq but also to produce new reference sequences for additional accessions of wheat or other species with highly complex genomes (Material s7).

The Lamp assembler provides capabilities to develop reference genome assemblies for various species and library types with reduced costs, offering flexibility in the design of an appropriate experimental workflow (Material s7). For separation of long reads mixed in a given super pool, Lamp offers a scalable and flexible ability to utilize short reads of each primary pool rather than sequencing barcodes. Moreover, in addition to the uses of the Lamp assembler for highly complex genomes, the system is generalizable for applications involving pooled samples sourced from multiple genomes of lower complexity. For example, a super pool may be produced by combining samples from dozens of bacterial strains or several fungal strains or from a mixture of BAC clones, viruses, bacteria, fungi, and plants in proportions corresponding to for the desired sequence coverage. In the case of small genomes, such as those of most fungal and bacterial accessions, Lamp is capable of completing a genome assembly using WGS data alone. As an example of a genome meeting these criteria of low complexity, the non-contiguous assembled genome of the dikaryon *Rhizoctonia cerealis* AG-DI strain consists of 16 pairs of chromosomes, in addition to a circular mitochondrial genome, with a total length of 83.4 Mb, and the various nuclei of the organism vary in their respective genomes to a low extent^60^. For slightly larger genomes, such as those of Arabidopsis and rice, Lamp can first use WGS data to produce contigs for a *de novo* reference assembly and then integrate data from BAC sequencing to fill remaining gaps. Finally, the Lamp assembler also offers improvements to reliability in assembling genomes from samples with a high level of contamination, for example, genomes from viruses, biotrophic bacteria and fungi, chloroplasts, mitochondria and low-copy plasmids, thus objectively reducing the labour and material costs needed to obtain samples of adequate volume and purity.

## Conclusion

In summary, we developed a low-cost genome assembly workflow based on a two-step pooling design and the Lamp assembler. Our application of this workflow significantly improved the continuity and accuracy of the 2DS of wheat in IWGSC RefSeq and furthermore established this genome portion as a benchmark for studying the assembly of complex genomes. The Lamp assembler itself is flexible and widely applicable for the assembly of genomes from diverse samples. By providing a means of assembling gold standard reference genomes with improved accuracy and reduced costs, our workflow accelerates the generation of reference genomes, in turn contributing to robust characterization of genomic variation and the resulting effects on traits.

## Materials and Methods

### MTP BAC library and bioinformatics data

The MTP BAC library TaaCsp2DSMTP from the *Triticum aestivum* cv. Chinese Spring chromosome arm 2DS was obtained from the French Plant Genomic Resources Centre, CNRGV (https://cnrgv.toulouse.inrae.fr/en/Library/Wheat). The MTP was assembled from the 2DS BAC library TaaCsp2DShA (Material s8) after WGP analysis^41^ and consists of 3,025 clones in eight 384-well plates (Table s1). The IWGSC RefSeq v1.0 wheat genome assembly and annotation, chromosome survey sequences^44^, WGP tags and 2DS physical map were accessed from the IWGSC sequence repository at the Unité de Recherches en Génomique Info (URGI, https://wheat-urgi.versailles.inra.fr/Seq-Repository). The Triticeae repeat sequence database (TREP) was accessed from GrainGenes (https://wheat.pw.usda.gov/).

### MTP BAC pooling and sequencing

The 2DS MTP clones were mixed into primary pools according to their positions on the 384-well plates received from CNRGV, with the exception of clones showing significant overlaps in the 2DS physical map, which were reassigned to simulate the fact that neighbouring clones in BAC libraries usually feature relatively little overlap. Each primary pool comprised 49-52 clones (recorded in Table s1). Prior to pooling, each clone was used individually to inoculate 13 mL of Luria-Bertani broth medium^61^ containing 12.5 μg/mL chloramphenicol in a 50 mL conical flask. Liquid cultures were incubated overnight at 37 °C and 225 rpm. Subsequently, the optical density (OD) at 600 nm of each culture was measured by a NanoDrop One^C^ (ThermoFisher, USA), and cultures were combined into primary pools in volumes calculated such that pools had an equal concentration of each clone.

BAC clones were extracted from each of these primary pools using either the Qiagen Large-construct Kit (10) (12462, Qiagen, Germany) or Omega BAC/PAC DNA Maxi Kit (D2154-02, Omega, China). For both kits, we followed the manufacturer’s standard instructions in the provided protocols. These protocols both begin with centrifugation of cells, resuspension of the pellet in a provided buffer and lysis of cells with an alkaline lysate solution. In the Qiagen protocol, isopropanol is added to the lysate to precipitate DNA, followed by an exonuclease treatment to digest chromosomal DNA. Next, the digested preparations are added to spin-columns, which are centrifuged to collect DNA in binding resin. The resin is washed by the addition of wash buffer to the resin and subsequent centrifugation, and finally, DNA is eluted by centrifugation of columns with elution buffer and collection of the eluent. The Omega protocol differs most notably in that DNA is collected by centrifugation without exonuclease treatment.

The resulting plasmid DNA samples were assayed using a Qubit^®^ Fluorometer 3 (ThermoFisher, USA) according to the protocol for the Qubit™ dsDNA HS Assay Kit (Q32851, ThermoFisher, USA). Each super pool was prepared by mixing up to 10 primary pools with respect to their concentrations to produce equal mixtures.

From each primary pool, an Illumina paired-end (PE) sequencing library was constructed with an average insert size of 350 bp. Libraries were sequenced using the Illumina HiSeq X Ten sequencing platform, producing approximately 15 Gb of 150-bp PE short reads. The throughput was further increased to as much as 40 Gb to ensure that approximately 15 Gb of short reads was produced from the BAC plasmids, with the remainder attributable to contaminants such as the *E. coli* chromosome. Library preparation and sequencing were performed by Novogene Co., Ltd.

For each super pool, a PacBio sequencing library was constructed with an average insert size of 10 kb or 20 kb. Long-read data sequencing was performed using the PacBio Sequel sequencing system. Library preparation and sequencing were performed at Tianjin Biochip Co., Ltd., or Novogene Co., Ltd. In cases where long reads of a primary pool totaled less than 300 Mb in size after being split in downstream methods as described below, additional long reads were further produced by appending the primary pool to another super pool.

### NODE contigs

PE short reads for each primary pool were end-trimmed to reduce sequences to high-confidence sequences of no more than 125 bp. Pairing was disregarded, and PE short reads were considered independently from one another for the purpose of constructing NODE contigs. To generate initial contigs from these reads, we used Velvet (version 1.2.10) with the parameter ‘*k*-mer length = 99’ to build *de Bruijn* graphs^45^. Subsequently, we disassembled all end-trimmed reads into 99-bp *k-*mers and built a collection of *k-*mers that were not included in initial contigs but had a coverage of 5× or more; these *k*-mers are referred to as “inter-contig *k*-mers.” As the end-trimmed reads had a maximum length of 125 bp, the disassembly of each produced up to 27 99-bp inter-contig *k*-mers. Initial contigs were also disassembled into 99*-*mers (termed intra-contig 99*-*mers), with their positions in the contigs noted. Finally, we built a hash table containing inter-contig 99*-*mers, intra-contig 99*-*mers, and the positions of the latter in initial contigs.

Next, we sought to build connections between 99-mers by identifying cases in which one of each originated from the same end-trimmed read. The hash table was searched according to the position order of 99*-*mers in each end-trimmed read to reveal 99-mer pairs overlapping in all but a terminal base for 98 bp of overlap. In cases where these pairs consist of both inter-contig and intra-contig 99-mers or two inter-contig 99-mers, a connection chain was recorded, with data including the sequences of each overlapping 99-mer, along with their relative orders, directions and overlaps.

To produce additional high-confidence connections, we constructed connection graphs by using initial contigs and inter-contig 99-mers as vertices and greedily extending inter-contig 99-mer vertices along both forward and reverse orientations using end-trimmed reads, accepting extensions meeting a threshold signal-to-noise ratio of 50:1. Extensions were terminated upon encountering a fork vertex, a dead end, an initial contig or a previously processed inter-contig 99-mer. Ultimately, the NODE contig set consisted of these extended contigs and initial contigs.

### Short-read- and paired-read-based connection chains

While NODE contigs are assembled using end-trimmed reads processed without regard to pairing and the trimmed portion, the construction of SCAF and COTG contigs requires connection chains built from paired reads. For each PE read pair, the correspondence of each read to NODE contigs was assessed by searching for matches in a hash table of 99-mers disassembled from NODE contigs, similar to the hash table process used during NODE contig construction. At this stage, reads found to match NODE contigs may either map to internal sequences of NODEs or to their edges; in the latter case, this extension leads to connection chains among NODEs, termed initial short-read-based connection chains (SR chains).

The lengths of gaps or overlaps between paired reads were estimated using knowledge of average library size and alignment of reads to NODE contigs. When both reads of a given pair mapped to the same NODE contig and the intervening NODE sequence was of a length typical of a gap in 350-bp insert libraries—between 175 bp (average insert size/2) and 525 bp (average insert size × 1.5)—the intervening NODE sequence length was taken to be the length of the gap or overlap. When two paired reads mapped to different NODE contigs, the length of the gap or overlap was calculated by subtraction of 225 bp (average insert size × 1.5 – read length × 2) from the lengths of the adjacent NODE regions matched by the paired reads. SR chains and NODE contigs represented by both reads in given read pairs were joined to form initial paired-read-based connection chains (PR chains), each featuring no more than a single gap.

Following the assembly of initial SR chains and initial PR chains, both chain types were used along with NODE contigs to form a connection graph. Prior to gap filling, this graph featured NODE contigs as vertices, linked only by initial SR chains. We attempted to fill each gap spanned by paired reads by using the left NODE of the gap as a starting point and seeking all extension paths composed of NODEs along the connection graph until the extension length exceeded the predicted gap size. NODE sub-chains sharing boundaries with a given gap were extracted to fill the gap in the initial PR chain, producing one or more filled PR chains. Paired reads with gaps remaining unfilled after this process were discarded. Next, we attempted to extend each NODE contig vertex in both forward and reverse orientations using a connection graph, accepting extensions meeting a signal-to-noise ratio threshold of 50:1. Each initial SR chain or filled PR chain was evaluated to determine whether all the internal connections were among these connection candidates; if not, then the chain was excluded from the final SR or PR chain set.

### COTG and SCAF contigs

Connection chains for COTG contigs were produced by cyclic extension of each NODE contig along forward and reverse orientations using SR chains. For each given NODE, a local connection graph was created using SR chains that contain NODE contigs. NODE contigs were extended along the local extension graphs. Upon encountering any fork in the graph, extension was continued using the top candidate if one met the 50:1 signal-to-noise ratio and 10× connection coverage thresholds. If such a candidate was found, it was appended to the existing connection chain or used to create a new one. When no single candidate meeting the specified criteria was found, extension was reattempted after rebuilding the local connection graph using SR chains matching terminal sub-chains consisting of multiple NODEs in proper order. Upon removal of duplications from extended chains, the final COTG contigs were produced.

Connection chains for SCAF contigs were produced by cyclic extension of PR chains along both orientations. For each attempt to extend a PR chain or its extension, we first evaluated whether the given chain could be extended along each orientation using SR chains as in the previously described COTG construction step. If a top candidate meeting the aforementioned thresholding criteria was identified, we next further detected all PR chains with perfect overlap to the boundaries by comparison of terminal sub-chains and selected the chain with the longest length for extension. Upon completion of extension for all PR chains, the extended chains were compared with one another to identify cases in which the chains were sub-chains of others. Upon removal of redundant chains, the final SCAF contigs were produced.

The SCAF contigs were scanned one by one to detect sub-sequences corresponding to COTG contigs or NODE contigs; these matches were recorded along with the sequence orientations and start and end positions. The correspondence of NODE contigs to COTG contigs was also detected in this manner. The SCAF, COTG and NODE contig sequences were merged to produce a final continuous contig set of long-read alignments; first, all SCAF sequences were imported, followed by the COTG sequences that were not sub-sequences of any SCAF sequences and, finally, NODE sequences that were not a portion of any SCAF or COTG sequences.

### Sequence alignments related to long reads

Contigs were aligned to long reads using the dual approach of pairwise alignment with BLAST (version 2.2.26)^62^ and multiple sequence alignment (MSA) with ClustalW^63^. Initial alignment was performed via BLAST using the parameters ‘*-p blastn -F F -v 1 -b 5000 -I T - G 2 -E 1 -q -1 -W 11*’. From these results, alignments meeting the thresholds of 70% identity and 150-bp alignment length were extracted. In cases in which two BLAST alignments meeting the aforementioned thresholds were extracted for a given long read, overlapping portions of the sequence alignments were compared to inform selection of the appropriate sequence. The identity score and length were recorded for each sequence alignment. In situations where a given alignment produced an identity score of 1.0% or an identity length that was five bp greater than that of any other alignment, it was selected for downstream assembly. Otherwise, portions of the long read and contigs were further compared by MSA.

ClustalW was used to align a given long read with multiple portions of contigs, using the parameters *‘-align -output=gde -case=upper -type=dna -pwgapopen=5.0 -pwgapext=1.0 - gapopen=5.0 -gapext=1.0 -gapdist=8 -maxdiv=40 -noweights’*. These parameters were also used for downstream alignment of long reads with one another during primary pool contig assembly (described below). All sequence alignments from MSA were stored as aligned base matrices in which each row represents a sequence and bases or gaps in sequences share a column in common when aligned. For each column in this matrix, matches between the long read and each given contig were counted. The alignment with the greatest number of matched loci was selected for assembly.

### Super-pool splitting and long-read-based connection chains

To ascertain the primary pools from which long reads in super pools were sourced, long reads of each super pool were aligned to contigs from all primary pools that had been pooled into the given super pool. When portions of a given long read were aligned with contigs from multiple primary pools, the alignments were investigated as follows to inform judgement of their source. The alignments were first compared, and any alignments that did not meet an overlap threshold of 300 bp were discarded. The mapping lengths of the retained alignments from each primary pool were then counted using the long read as the coordinate axis. Each long read was assigned to the primary pool with the maximum mapping length, as well as any primary pools with mapping lengths at least 150 bases less than the maximum mapping length. Long reads could be assigned to two or more primary pools in certain situations, particularly if BAC clones with overlaps were pooled into adjacent primary pools.

To map NODE contigs to the long reads, final contigs for every primary pool were first aligned to each long read. All alignments of long reads to SCAF contigs were subsequently downgraded to alignments to corresponding COTG or NODE contigs on the basis of COTG or NODE orientations and positions within SCAF contigs, producing two or more COTG or NODE alignments from a given SCAF alignment. To inform selection of the most appropriate alignment, alignments showing overlaps of at least 114 bp (*k*-mer length × 1.15) in length were compared using base identity as a basis to select the ideal NODE alignment or aligned portion of a COTG alignment. Unselected alignments were discarded. The COTG alignment was completely excluded in situations in which the retained alignment was less than 198 bp (*k*-mer length × 2) in length. Once all alignments were compared and filtered, the retained COTG alignments or retained COTG portions were further downgraded to NODE alignments. Redundancy of NODE alignments was removed from the results, and alignments were removed if at least one unaligned end of the aligned NODE contig or long read exceeded 30 bp in length. Subsequently, alignments were removed from the results if their estimated length of overlap with other alignments exceeded 114 bp (*k*-mer length × 1.15). The retained alignment results were then converted to raw long-read-based (LR) connection chains of NODE contigs.

The raw LR chains were polished to revise inaccurate, gapped or otherwise faulty internal connections. For each connection with an estimated length of overlap between 79 bp (*k*-mer length × 0.8) and 119 bp (*k*-mer length × 1.2), the overlap size was re-evaluated by alignment to SR and PR chains and revised to reflect the overlap length indicated by these higher-confidence sequences. For any connections with an overlap size exceeding 98 bp (*k*-mer length - 1) or with any surrounding NODE contig of below 30× *k*-mer coverage, all surrounding NODE contigs were temporarily removed, leaving a new gapped connection. For all gapped connections, gap filling was attempted as previously described for the gap filling step of initial PR chain construction. If this process yielded no candidate to fill the gap, the connection was reset to the previous state. For any gap with two or more candidates, MSA was performed, and differential loci between candidates and long reads were tallied. The candidate presenting the fewest differential loci was deemed the ideal choice and retained for downstream assembly.

### Primary pool contig assembly

The process of producing contigs for each primary pool began with the initialization of seeds, which were NODE contigs or chains constructed from gapless matrices of NODE contigs. A given seed was then extended in both orientations using LR chains that contained the seed. In accordance with the appropriate direction for extension, columns in the aligned base matrix were scanned one by one to reveal potential extension paths. Candidates for extension were accepted when they met the 10:1 signal-to-noise ratio and 10× connection coverage thresholds. Upon encountering a fork with multiple candidates for extension, the scan was suspended, and the corresponding LR chains were manually reviewed to inform the decision for extension.

When generating the sequence of genome chains, the paired NODE contigs of candidate connections in the chain were joined to form a single continuous sequence if the overlaps were exactly the same; otherwise, a gap was retained. For a given gap, NODE contigs connected through the chain over a distance of up to 10 kb were scanned, and their occurrence within all genome chains in the primary pool was tallied. For each non-repetitive NODE contig, long reads mapped to the same LR chains as the given NODE contig were collected, and sub-sequences corresponding to the gap were extracted along with flanking sequences of at most 1,000 bp. MSA was performed on extracted sequences, and consensus sequences were built from the resulting aligned base matrix. The best choice of all consensus sequences was determined by MSA and used to close the gap.

### Chromosome-scale contig assembly and chromosome anchoring

Chromosome-scale contigs were greedily assembled by utilizing overlaps among BAC sequences. The overlaps for a given BAC sequence were initially identified by aligning the 5 kb terminal sequences of each BAC sequence against those of all others using BLAST with the parameters *‘-p blastn -FF -G2 -E1 -e 1e-4’*. The candidates were first filtered by a threshold of 99% identity and were evaluated by manual review of dot-plot visualization results generated using the dotmatcher tool in EMBOSS version 6.5.7.0 with the parameters *‘-windowsize 100 -threshold 90’*.

Each chromosome-scale contig was aligned to all survey sequence sets specific to whole chromosomes or chromosome arms using BLAST with the parameters *‘-p blastn -FF -v1 -b 10000 -e 1e-10*’. The total length of matching bases in each survey sequence set was determined, with alignments accepted if they met the 300-bp matched length threshold and shared 100% identity. Contigs were then anchored to the IWGSC RefSeq v1.0 using the same BLAST parameters and thresholding criteria used when detecting overlaps between BAC sequences during chromosome-scale contig assembly. Large structural differences were identified by manual inspection of the dot-plot visualizations made with MacVector version 17.0.3 (https://macvector.com/).

### Aligning BAC sequences to the corresponding BAC clones

To match BAC sequences from primary pools to their BAC clones of origin, BAC sequences from each primary pool were cross-referenced with all WGP tags for each BAC clone. For each primary pool, exact matches to each whole WGP tag were tallied. When any BAC sequence from a primary pool matched with less than 80% of WGP tags, with only one or two WGP tags, or matched two or more clones, these results were manually inspected to remove false positives. For primary pool BAC sequences that were unmatched with their clone of origin, the search was extended to the MTP library and the source library, and the results were retained when 80% or more of these tags were perfectly retrieved.

### Annotation

Gene models were predicted by the BRAKER pipeline (version 2.1.4) using training sets generated from the high-confidence (HC) or low-confidence (LC) protein models of IWGSC RefSeq v1.0 and RNAseq data^64^. The predicted gene models were integrated by EVidenceModeler (v1.1.1)^65^. Functional annotation of these gene models was performed using eggNOG (Emapper-2.0.1 emapper DB: 2.0)^66^.

### Primer design, PCR amplification and Sanger sequencing

Repeat junctions in non-2DS contigs, as well as large mis-assemblies in 2DS, were identified by searching the TREP database by BLAST and inspecting results to identify junction positions^48^. Primers for amplification of repeat junctions were then designed using Primer3 (version 2.4.0) with the desired amplicon size set to range from 150 to 650 bp^67^. PCR was then performed using these primers with 2× Taq DNA Polymerase Master Mix (Vazyme, China) and the following thermocycler configuration: denaturation at 94 °C for three min, followed by 32 cycles of denaturation at 94 °C for 30 s, annealing at 55-60 °C for 30 s, and extension at 72 °C for 40 s, and a post-PCR final extension step at 72 °C for 10 min. PCR products were then separated by electrophoresis on 0.8% agarose gels.

For long-range amplification of sequences encompassing boundaries of 2DS large mis-assemblies, primers were designed to amplify sequences ranging in length from 1,500 to 7,500 bp. PCRs with KOD FX DNA polymerase (KFX-101, Toyobo Co., Ltd.) were performed as follows: denaturation at 94 °C for 2 min, followed by 36 cycles of denaturation at 98 °C for 10 s, annealing at 60 °C for 30 s, and extension at 68 °C for 5 min. The resulting PCR products were then separated by electrophoresis on 1% agarose gels. Samples were prepared for Sanger sequencing at concentrations based on their respective amplicon sizes predicted by Primer3 and submitted to TsingKe Co., Ltd. Finally, the Sanger sequences were subjected to *in silico* manual assembly, performed using MacVector.

---

**Supplemental Figure 1.**
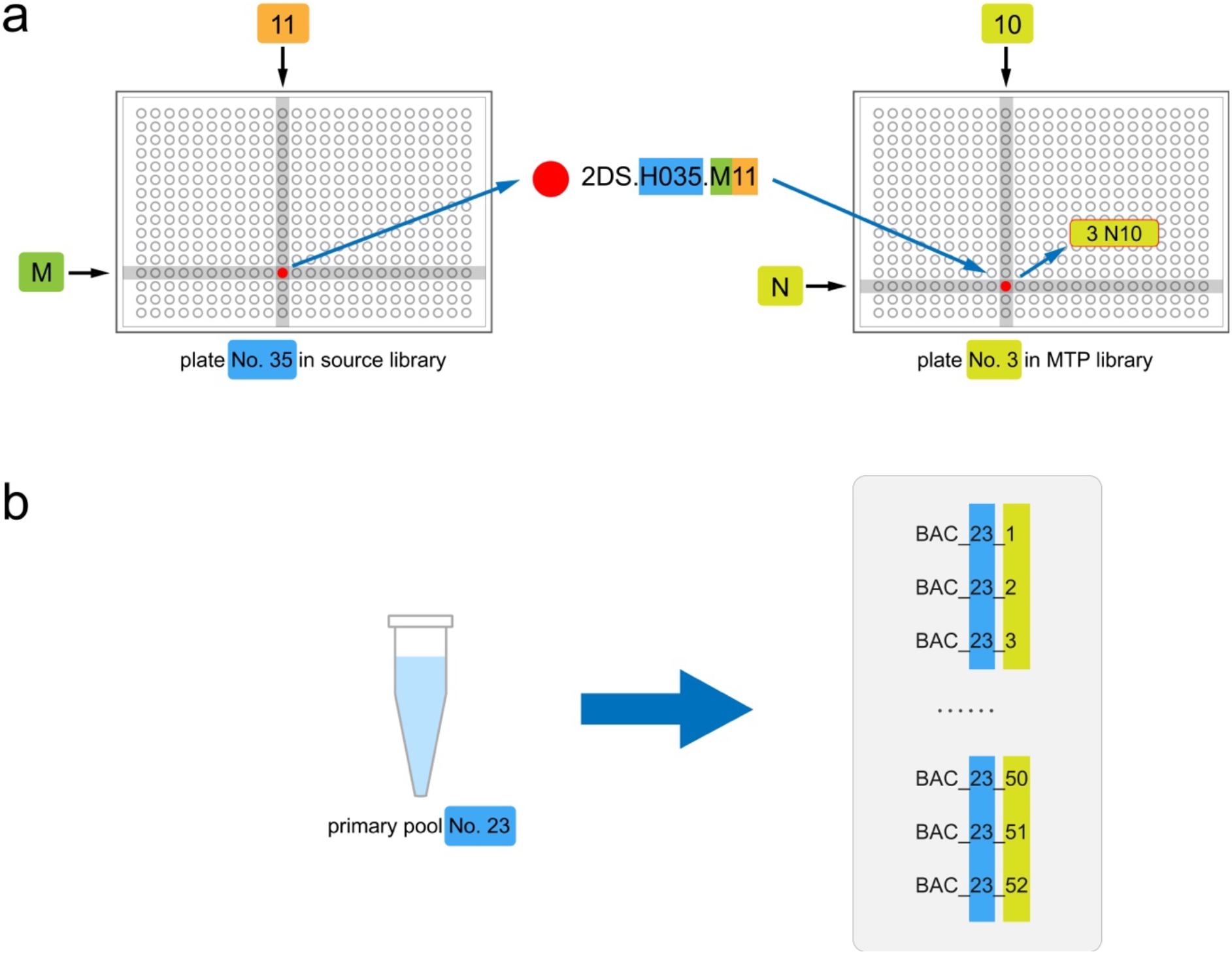
Nomenclature of BAC clones and BAC sequences: **(a)** The BAC clone was named according to its well position in the source library. In the MTP library, its well position was recorded separately. **(b)** The BAC sequence was named according to the number of primary pools, followed by a number assigned during sequence assembly.

**Supplemental Figure 2.**
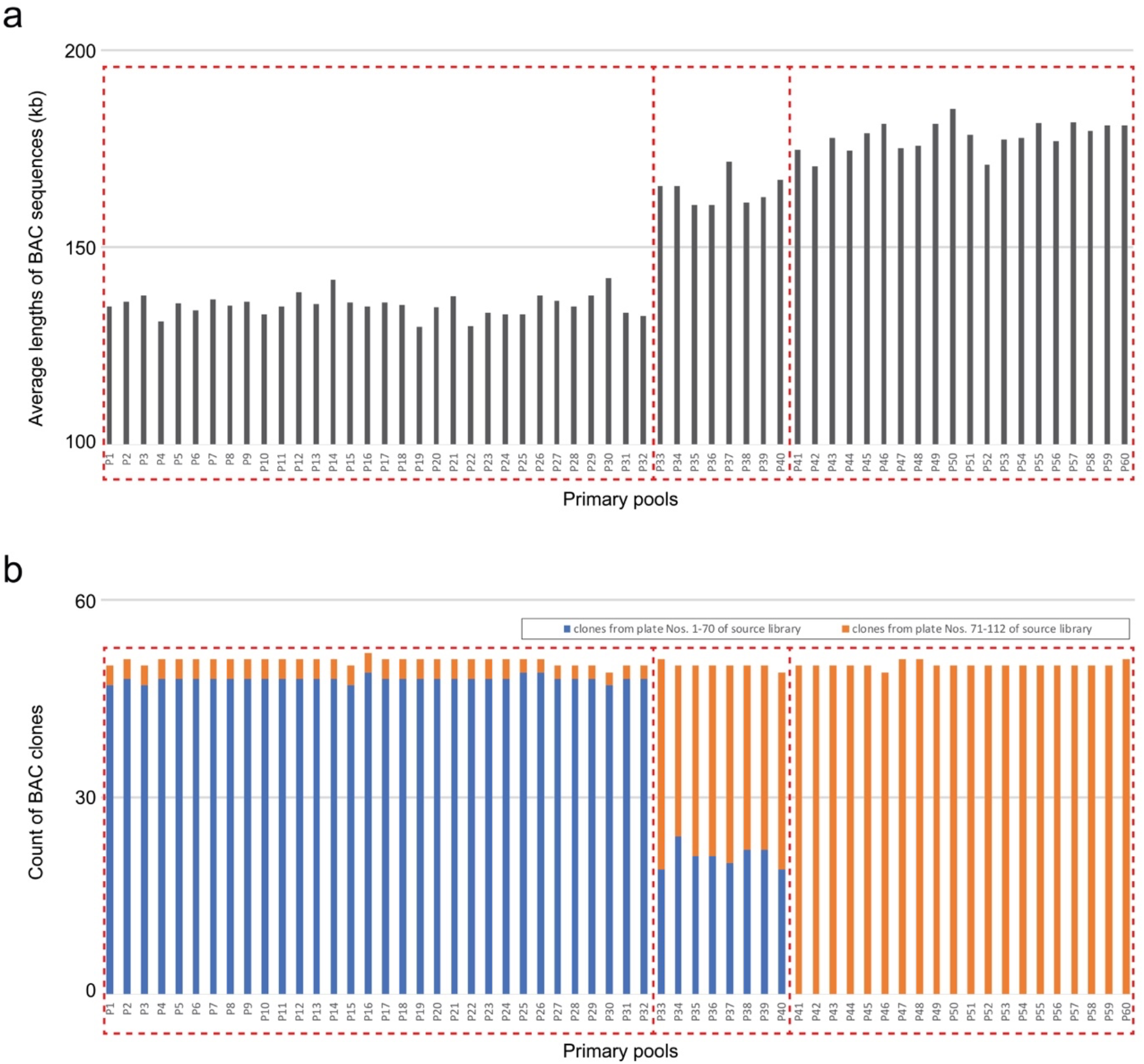
Bar charts showing that insert lengths differed among the 60 primary pools, which is consistent with the 2DS source library in which clones in plate Nos. 71-112 were detected to have a greater average insert size than those in plate Nos. 1-70 (*Isolation of BAC DNA and insert analysis* section in Material s8). In our BAC assembly, the insert lengths range between 130-142 kb for primary pool Nos. 1-32, 171-184 kb for pool Nos. 41-60, and 161-172 kb for pool Nos. 33-40 **(a)**, corresponding to clones in source library plate Nos. 1-70 and Nos. 71-112 and a mixture of the above two fractions, respectively **(b)**.

**Supplemental Figure 3.**
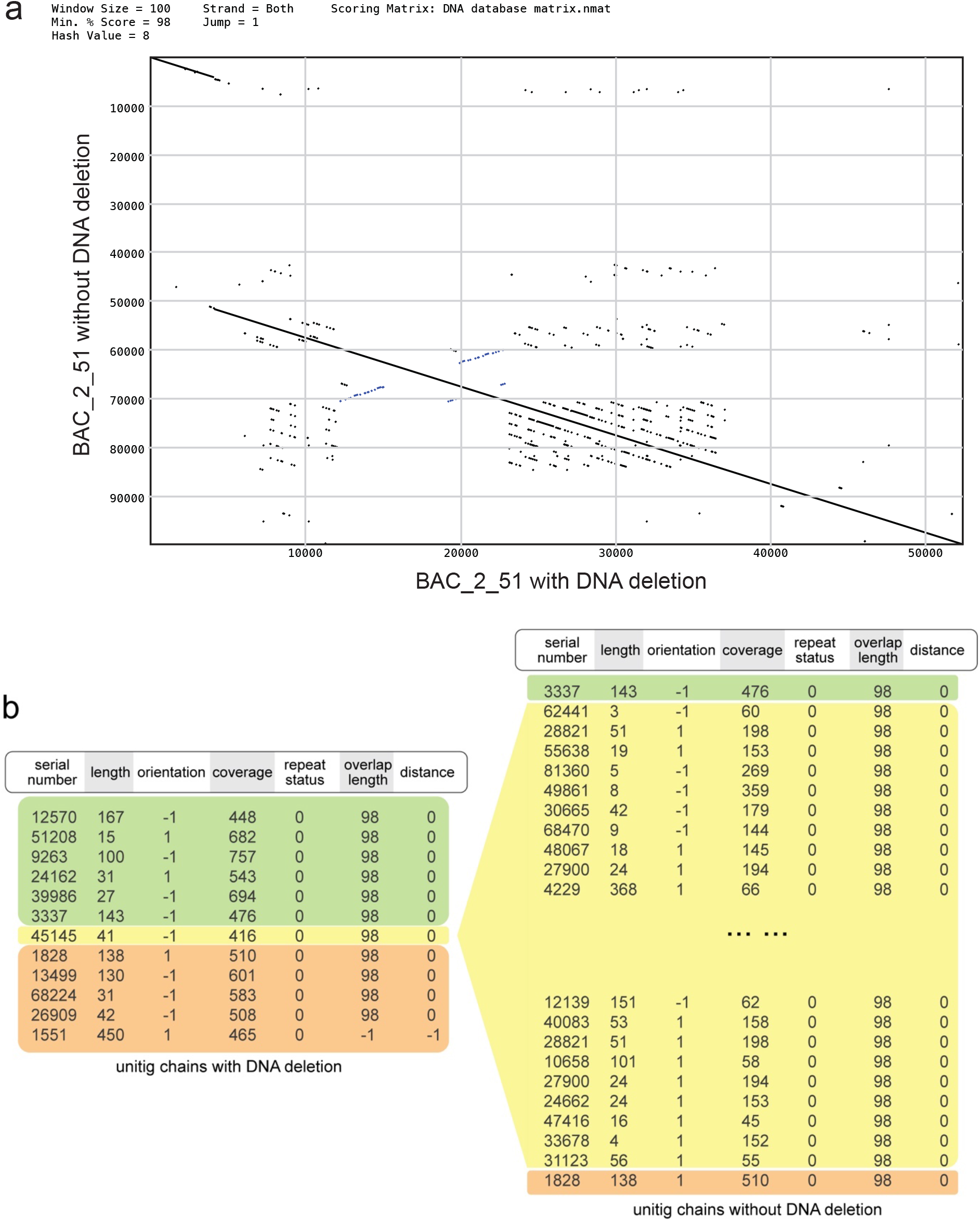
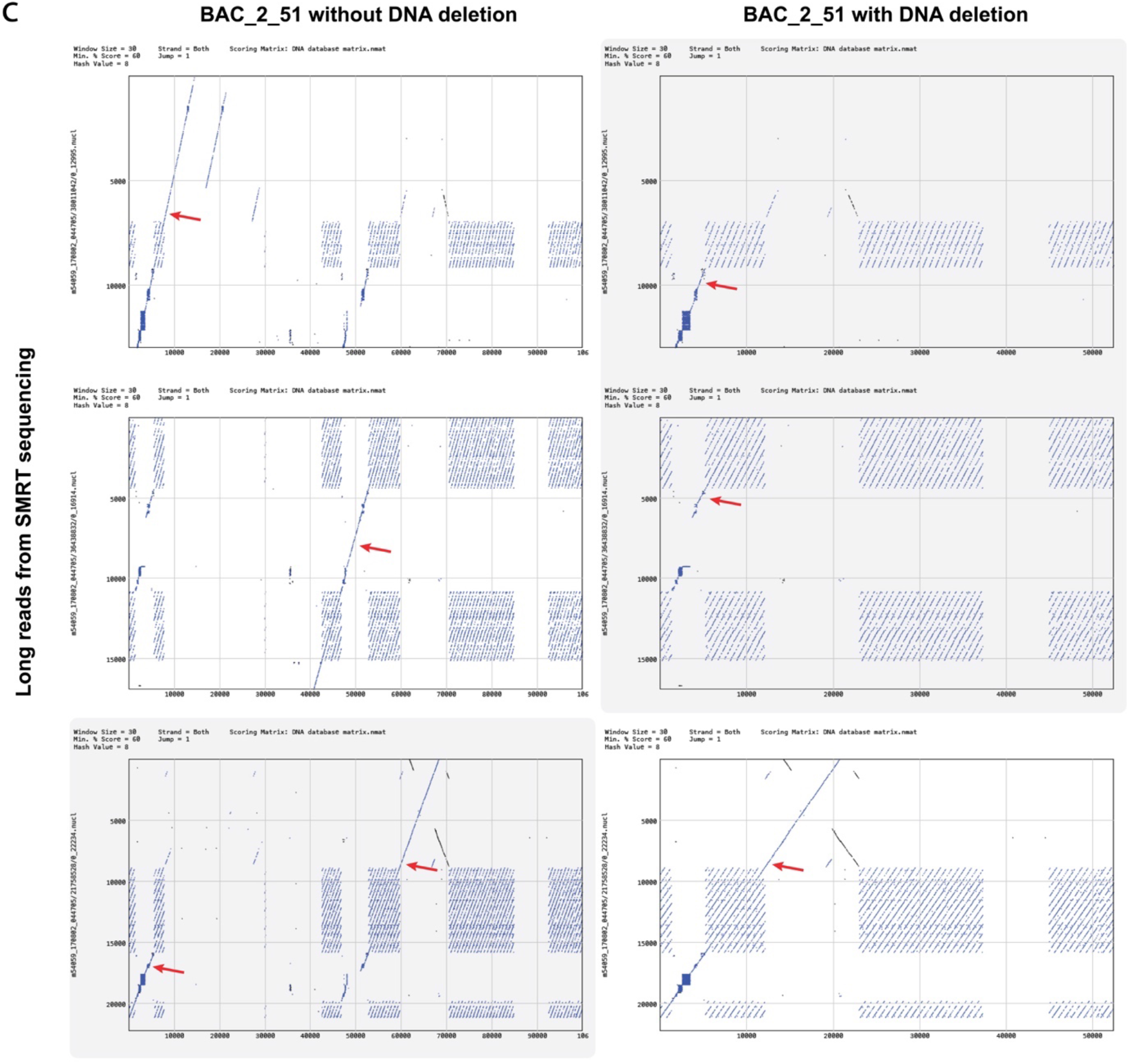
Illustration of a putative plasmid DNA deletion event in the clone with the sequence of BAC_2_51: **(a)** Dot-plot visualizations of assembled BAC sequences with and without the putative deletion event. **(b)** The short-read-based unitig chain of BAC_2_51 forked in the *de Bruijn* graph is shown. Each row represents a unitig in the chain, for which attributes (stored as tab-delimited text) include serial number, *k*-mer based length, orientation, coverage, repeat status (a placeholder that is reserved for use in the future; its values were all set to 0 here), overlap length and distances to adjacent unitigs. **(c)** The dot-plot visualizations of long reads support both assemblies (with and without DNA deletion). The first and second rows show dot-plot visualizations for long reads that support the assembly without DNA deletion, whereas the third row provides long-read-based evidence for the putative plasmid DNA deletion event. Red arrows mark lines showing collinearity between BAC sequences and long reads.

**Supplemental Figure 4.**
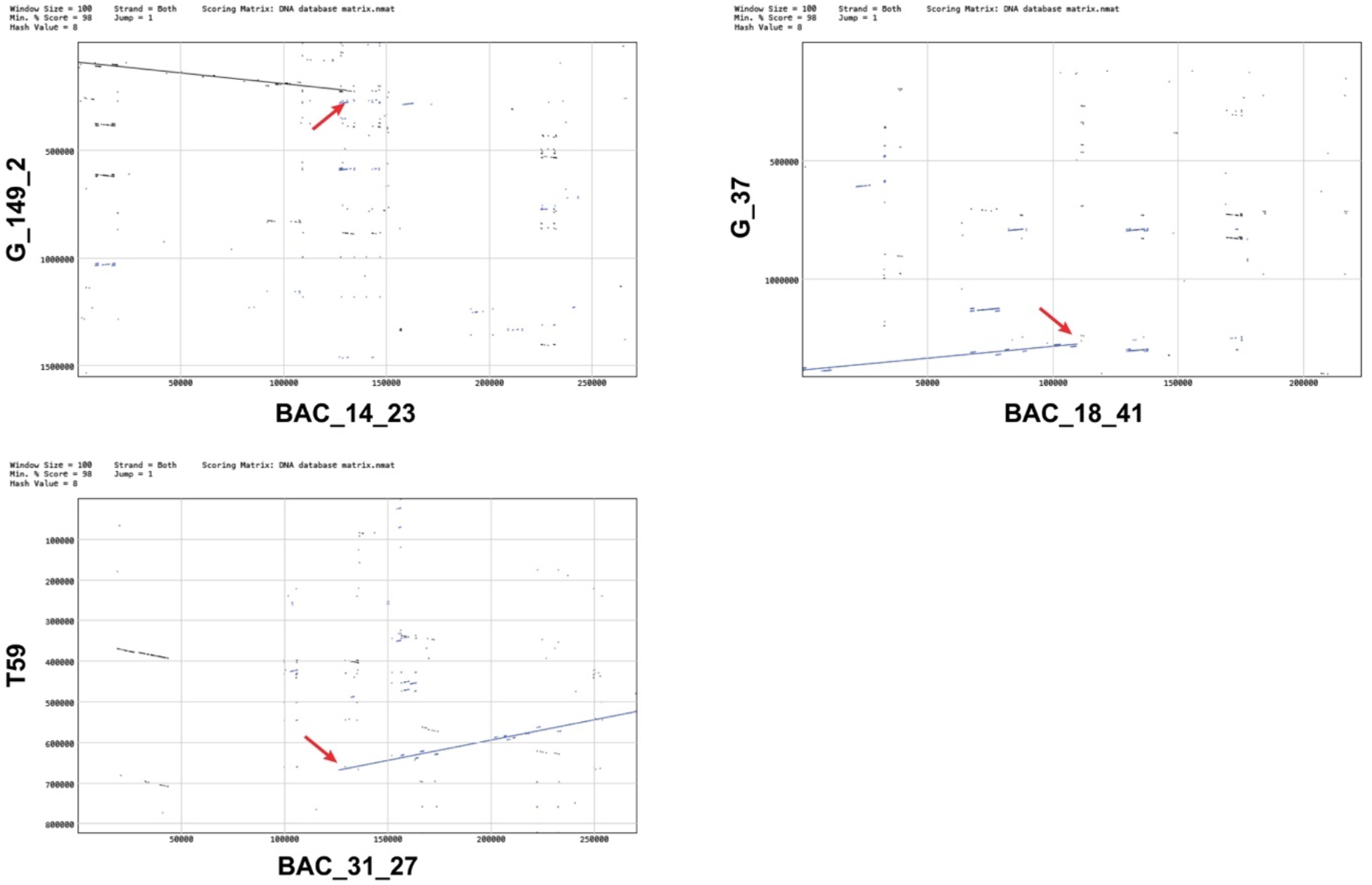
Dot-plot visualizations for alignment of chimeric BAC sequences against contig sequences with partial overlaps. Red arrows mark the junction point (x-axis) in chimeric sequences.

**Supplemental Figure 5.**
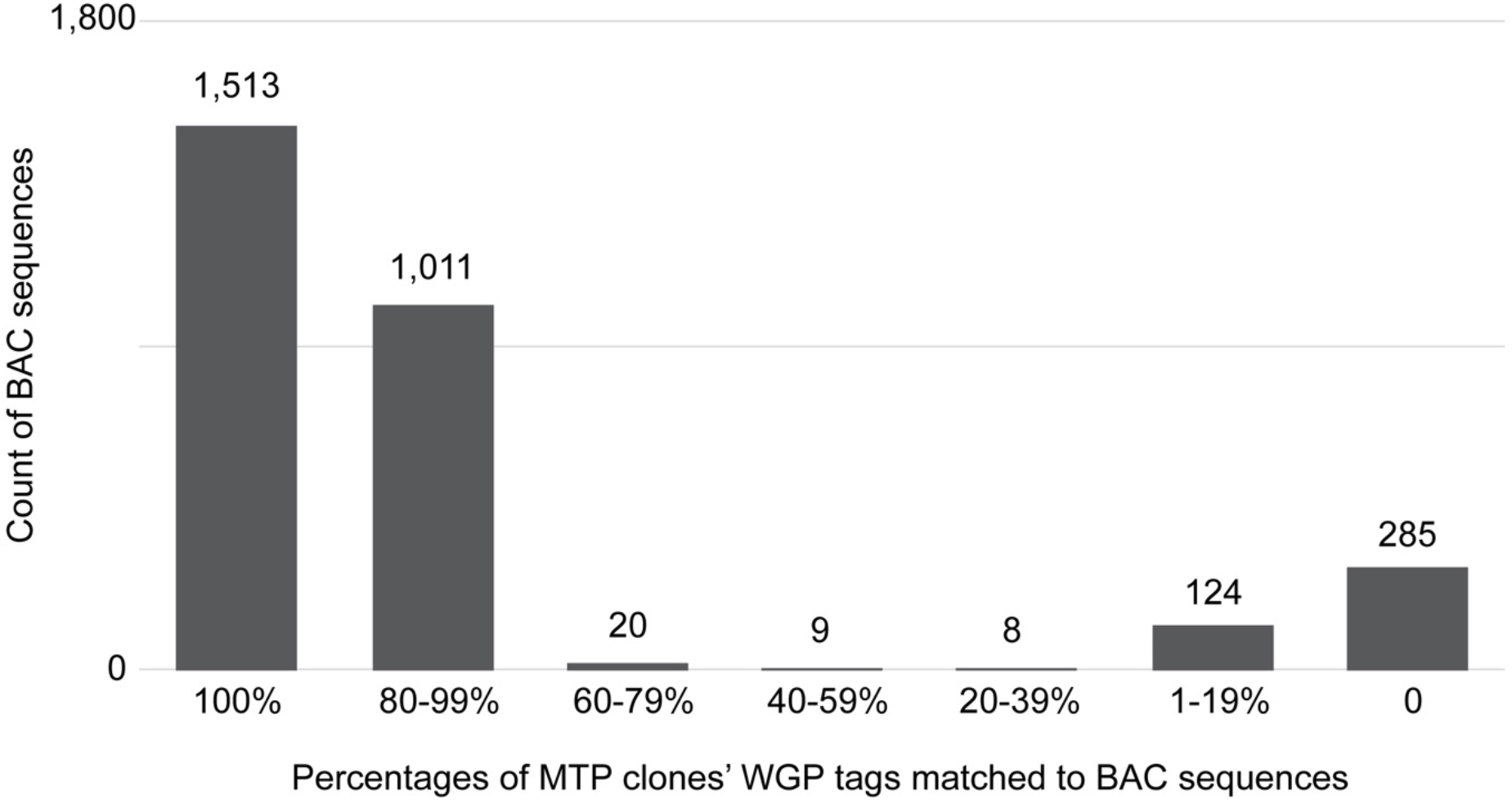
Proportional distribution of MTP clones’ WGP tags matched to BAC sequences.

**Supplemental Figure 6.**
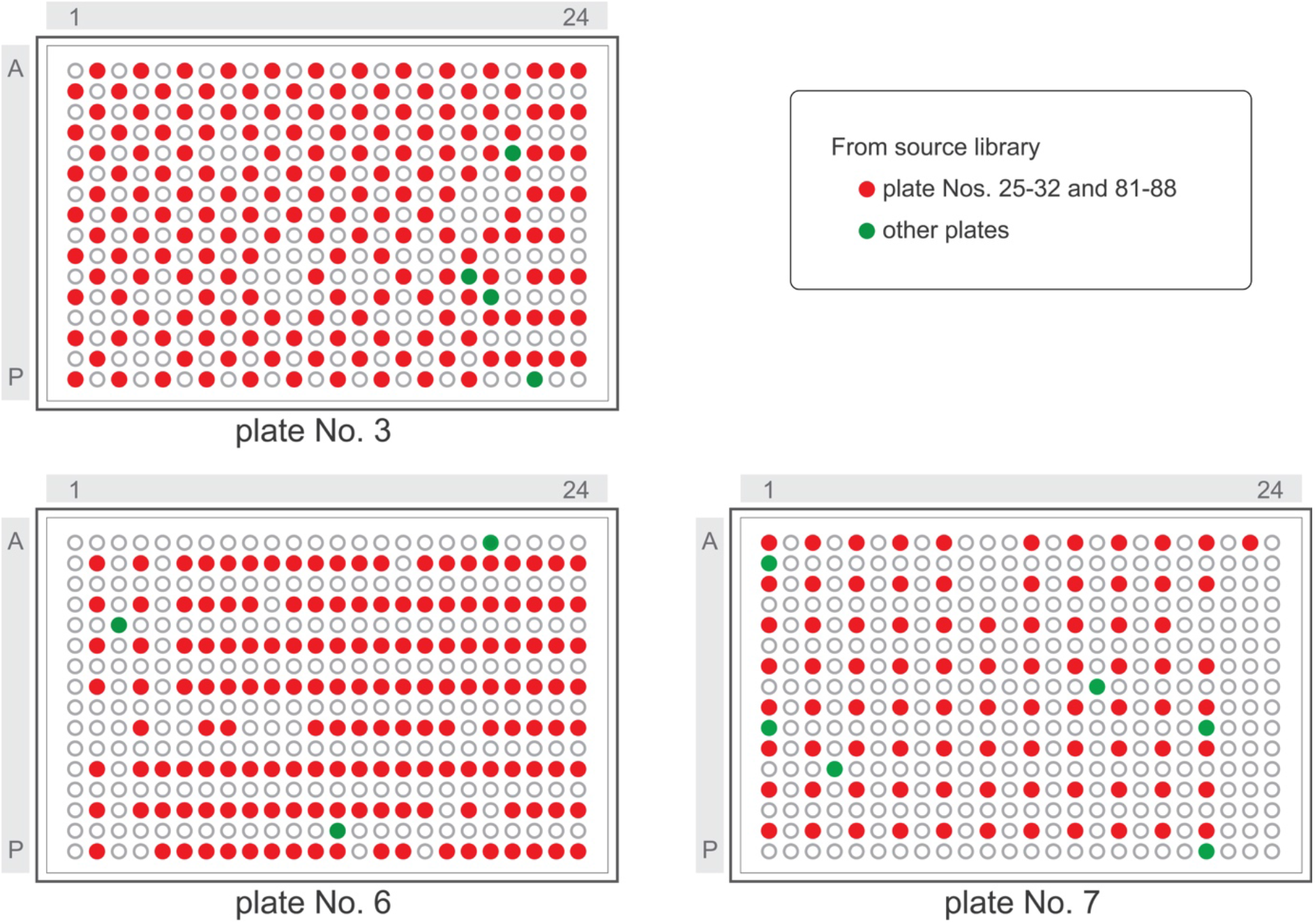
Well positions of unmatched BAC clones in 384-well plate Nos. 3, 6 and 7 of the MTP library received from CNRGV.

**Supplemental Figure 7.**
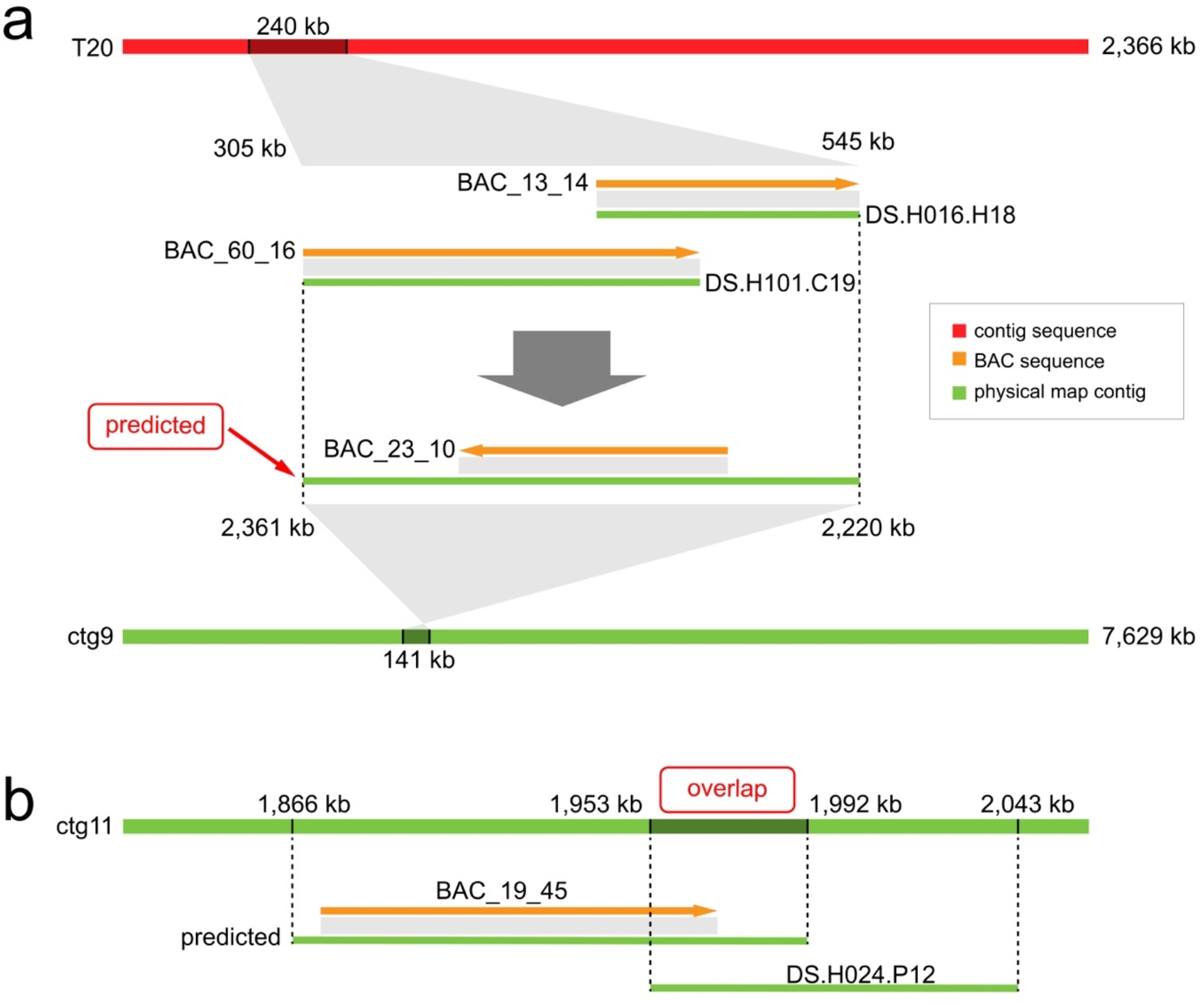
Illustration of a workflow to re-anchor BAC sequences to a physical map and detect their potential overlaps with the unmatched MTP clones: **(a)** The BAC_23_10 sequence was re-anchored to the ctg9 physical map contig, with an estimated position in the physical map ranging from 2,220-2,361 kb based on its adjacent BAC sequences on the T20 contig. **(b)** An overlap candidate on the ctg11 physical map contig was detected between the BAC_19_45 sequence and DS.H024.P12 clone.

**Supplemental Figure 8.**
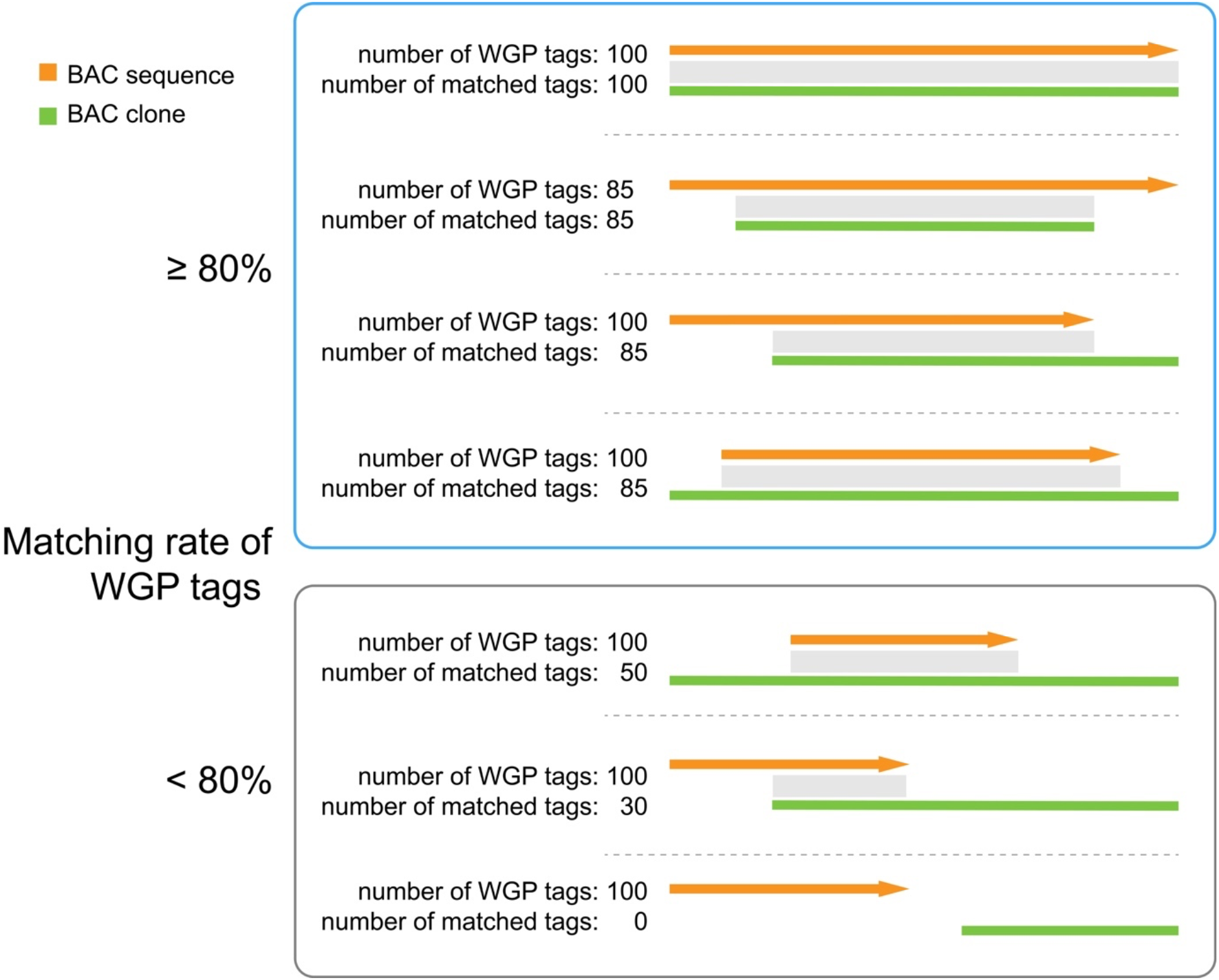
Illustration of how one BAC sequence might match multiple BAC clones in the MTP library or source library, as shown in Tables s12, s13 and s14.

**Supplemental Figure 9.**
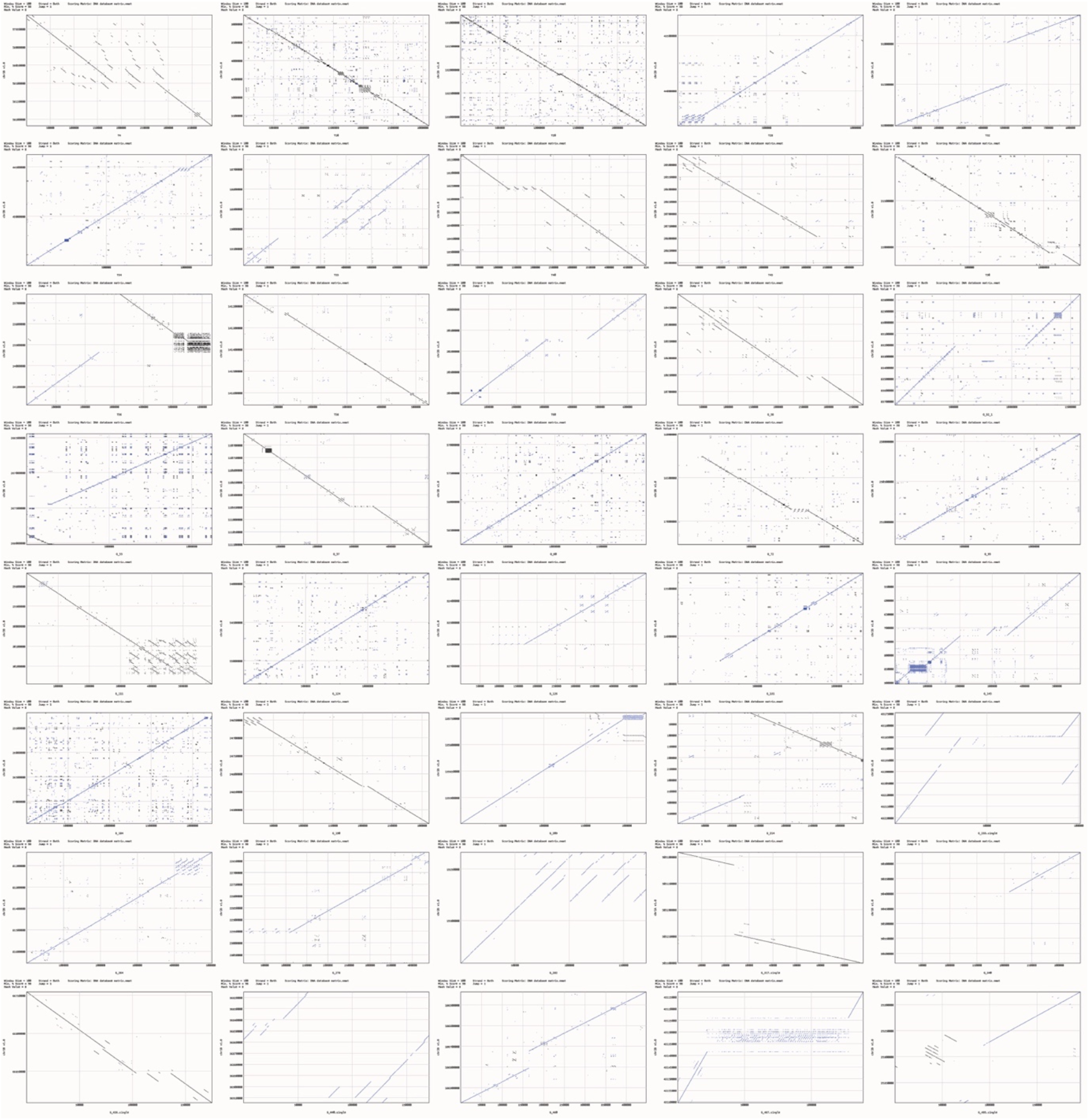
Dot-plot comparisons of contig sequences aligned against corresponding sequences in IWGSC RefSeq v1.0 reveal large structural differences.

**Supplemental Figure 10.**
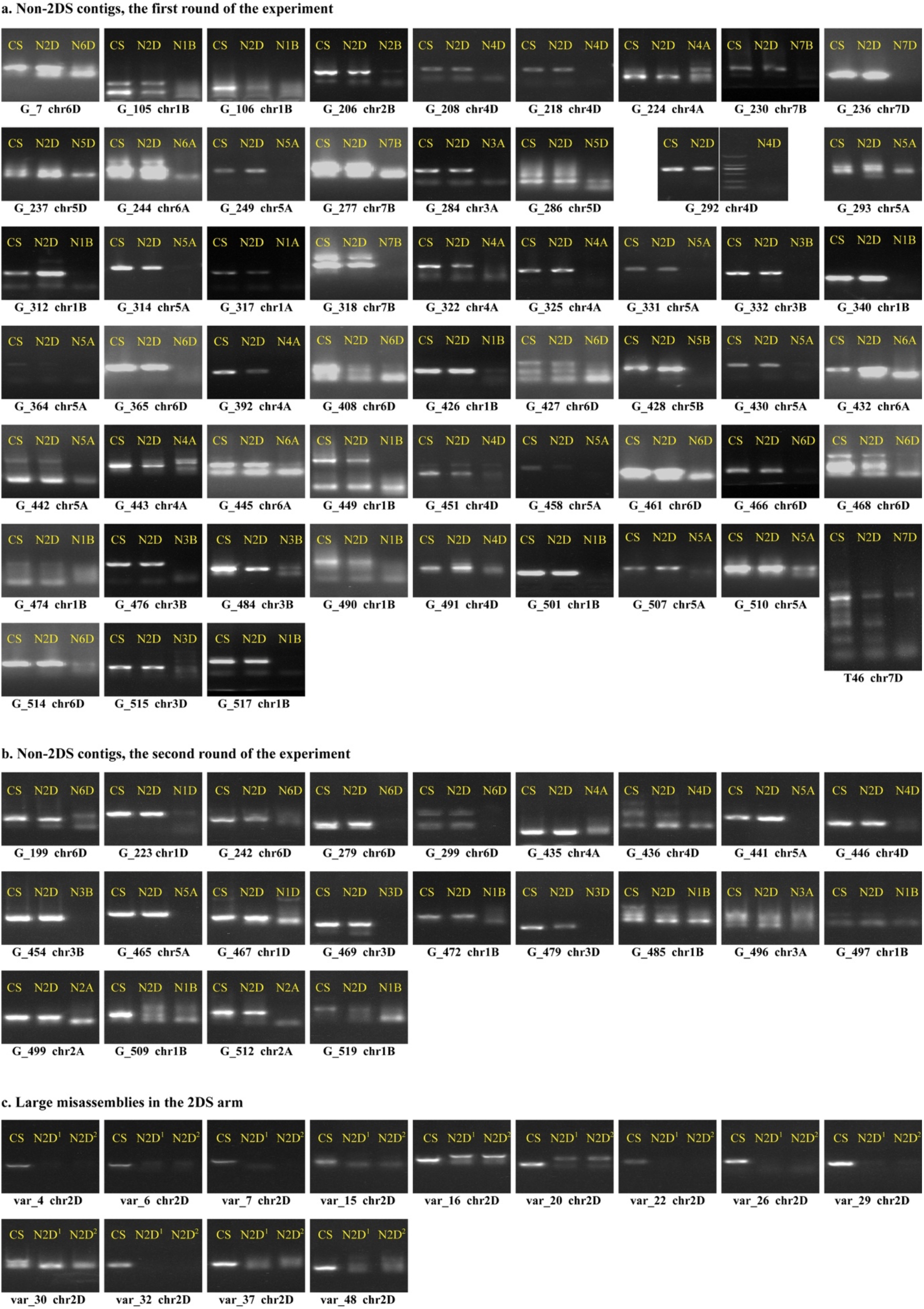
The results from chromosome anchoring experiments confirm chromosomes of origin for non-2DS contigs **(a, b)** and large misassemblies in 2DS **(c)**. DNA from nullisomic-tetrasomic lines of Chinese Spring wheat were used as templates for PCR amplification in all of these experiments. Electrophoresis of PCR products generally revealed amplicons of the expected sizes. For these non-2DS contig anchoring tests, positive bands of approximately equal size were amplified from both wild-type Chinese Spring and the N2D line, while absent or polymorphic bands resulted from amplification using template DNA from the predicted nullisomic-tetrasomic line for the corresponding chromosome. For large misassemblies in the 2DS, positive bands were detected only for the wild-type Chinese Spring and not for the nullisomic-tetrasomic line templates N2DT2A (N2D^1^) and N2DT2B (N2D^2^).

**Supplemental Figure 11.**
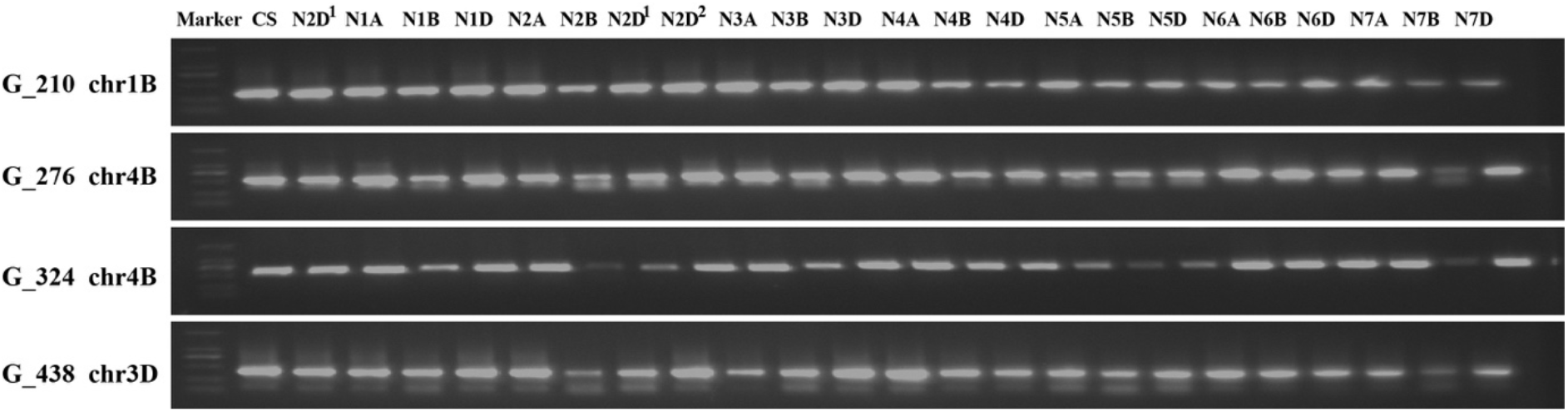
Additional chromosome anchoring experiments were performed using a full set of nullisomic-tetrasomic lines and four primer pairs randomly selected among non-2DS contigs indicated by previous experiments (Figure s10) to be unanchored. Wild-type Chinese Spring was used as a control. Notably, these experiments do not indicate anchoring to any chromosome, suggesting that the unanchored contigs or large misassemblies were not anchored due to nonspecific primers.

**Supplemental Figure 12.**
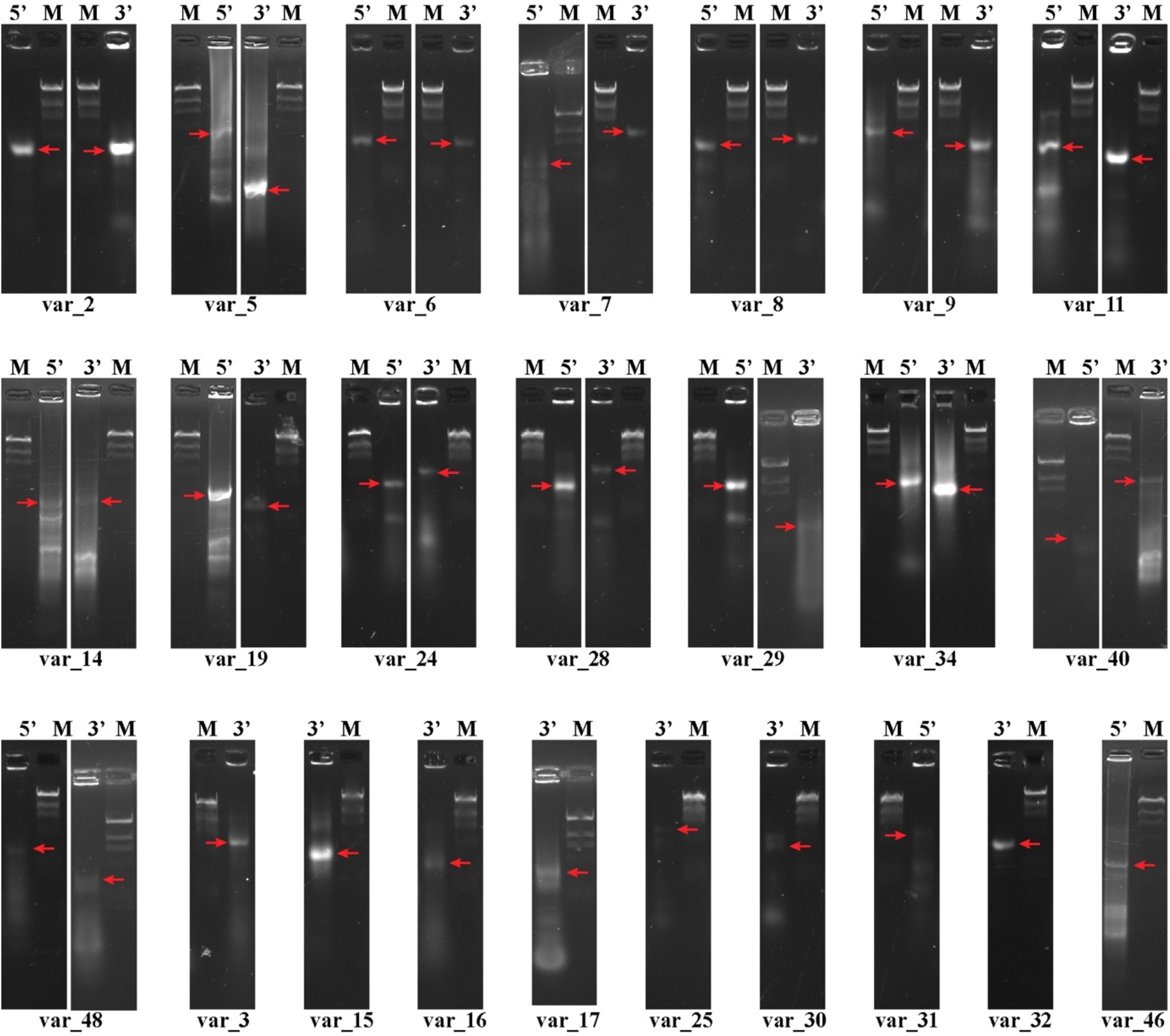
Long-range amplification results surrounding the boundaries of 2DS large misassemblies by using a DNA sample from Chinese Spring as a template. The PCR products were evaluated by electrophoresis analysis. The red arrows indicate the bands with the same size as those predicted from our assembly.

**Supplemental Figure 13.**
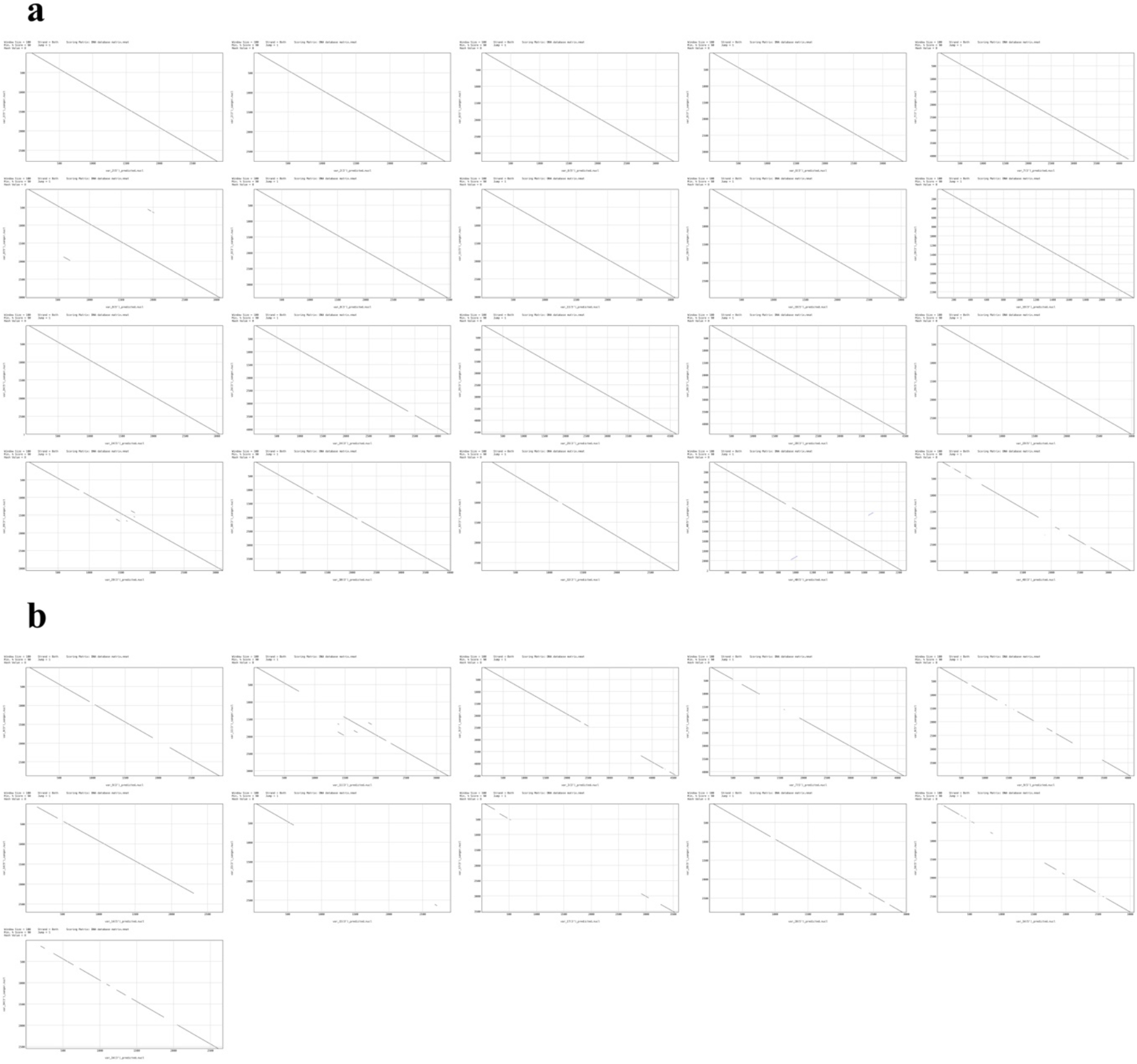
Dot-plot visualizations for Sanger sequencing results of target bands from long-range amplification (Figure s12) to the products predicted from our assembly: **(a)** Comparison of bands that were sequenced successfully; **(b)** comparison of bands with some portions not sequenced successfully.

**Supplemental Table 1. Pooling design of the TaaCsp2DSMTP BAC library:** The coordinates of MTP clones in 384-well plates as received from CNRGV are shown along with the positions of these clones in physical map contigs. The nomenclature of BAC clones is shown in Figure s1.

**Supplemental Table 2. Characteristics of BAC sequences:** All of these data were calculated for each BAC sequence following end-trimming of vector sequences.

**Supplemental Table 3.** Average insert sizes of BAC clones in primary pools.

**Supplemental Table 4.** Characteristics of contig sequences.

**Supplemental Table 5. Positions and orientations of BAC sequences in contig sequences.**

**Supplemental Table 6. Summary of portions in contig sequences that align with genome survey sequences from each chromosome or arm:** These summary statistics were produced for alignments with 100% identity over a minimum length of 300 bp. The abbreviations for each genome survey sequence set are listed in the footnote.

**Supplemental Table 7. Details of contig sequences aligned to genome survey sequences of each chromosome or arm:** Details are listed for all alignments with 100% identity over a minimum length of 300 bp. The abbreviations for each genome survey sequence set are the same as those listed in the footnote of Table s6.

**Supplemental Table 8.** Details of the results from anchoring contig sequences to IWGSC RefSeq v1.0.

**Supplemental Table 9. Correspondence between BAC sequences and MTP clones.**

**Supplemental Table 10. Details of MTP clones matching multiple BAC sequences from the same primary pool.**

**Supplemental Table 11.** Details of positional comparisons in the physical map between unmatched BAC sequences and unmatched clones from plate Nos. 25-32 and 81-88 of the source library: Following the workflow illustrated in Figure s7, the positions of unmatched BAC sequences in the physical map were estimated using adjacent BAC sequences in the contig sequences listed in Table s5. Marked in red letters are the detected positional overlaps of the BAC sequence to specific clones from the same primary pool.

**Supplemental Table 12.** Summary of alignments between unmatched BAC sequences and BAC clones with WGP tags.

**Supplemental Table 13. Alignments of unmatched BAC sequences to clones with WGP tags in the MTP library:** A given BAC clone was considered to be aligned to an MTP clone if at least 80% of WGP tags for the clones were detected in the BAC sequence.

**Supplemental Table 14. Alignments of unmatched BAC sequences to MTP clones with WGP tags in the source library:** A BAC clone was considered to be aligned based on the same criteria used in Table s13.

**Supplemental Table 15. Correspondences between contig sequences and physical map contigs:** Contig sequences containing at least five MTP clones corresponding to unique BAC sequences are listed. Marked in red letters are contig sequences matching two or more portions of physical map contigs.

**Supplemental Table 16. Positional relationships between physical map contigs and BAC sequences in contig sequences:** Five chimeric BAC sequences and 16 BAC sequences listed in Table s10 are not included.

**Supplemental Table 17.** Alignments of physical map contigs and IWGSC RefSeq v1.0 using the assembled contig sequences as a medium.

**Supplemental Table 18. Characteristics of large structural differences between contig sequences and IWGSC RefSeq v1.0.**

**Supplemental Table 19.** Functional annotations of large structural differences.

**Supplemental Table 20.** Summary of large structural differences matched to the unanchored sequences (chrUn) in IWGSC RefSeq v1.0: Summary statistics are shown for alignments of 99% identity and greater over lengths of at least 3 kb.

**Supplemental Table 21.** Details of large structural differences matching to the unanchored sequences (chrUn) sequence in IWGSC RefSeq v1.0: Details are provided for the same alignments as featured in Table S20, specifically for alignments with 99% identity and greater over lengths of at least 3 kb.

**Supplemental Table 22.** Summary of portions of large structural differences aligned to genome survey sequences of each chromosome or arm: Summary statistics are shown only for alignments of 100% identity over a minimum length of 300 bp. Genome survey sequences are labelled with abbreviations as in Table s6.

**Supplemental Table 23.** Details of large structural differences aligned to genome survey sequences of each chromosome or arm: Details are provided for the same alignments as in Table s18, with 100% identity over a minimum length of 300 bp. Genome survey sequences are labelled with abbreviations as in Table s6.

**Supplemental Table 24.** Primers used in chromosome anchoring experiments for non-2DS contigs and 2DS large misassemblies.

**Supplemental Table 25.** Primers used for PCR amplification and Sanger sequencing to investigate and confirm boundaries of 2DS large misassemblies.

**Supplemental Material 1.** The NODE unitigs of each primary pool are provided in FASTA format.

**Supplemental Material 2.** Genome connection chains for BAC sequences.

The files from ‘P1.txt’ to ‘P60.txt’ correspond to the 60 primary pools. Each file contains genome connection chains of all BAC sequences from a given primary pool. Genome chains are represented in a format similar to FASTA, with a description line such as ‘>BAC_1_1’ that states the name of the BAC sequence, followed by tab-delimited lines to define attributes of the connected unitigs, including serial number, *k*-mer based length, orientation, coverage, repeat status (a placeholder reserved for use in the future), overlap length and distance to adjacent unitigs.

**Supplemental Material 3.** The assembled BAC sequences are provided in FASTA format.

**Supplemental Material 4.** The assembled chromosome-scale contig sequences are provided in FASTA format.

**Supplemental Material 5.** Gene model sequences are provided in FASTA format.

**Supplemental Material 6.** Summary of comparison to IWGSC RefSeq v2.0.

RefSeq has been improved with the v2.0 update, which utilizes WGS PacBio reads and genome optical mapping (http://www.wheatgenome.org/). In RefSeq v2.0, a large number of misassemblies and gaps were revised, resulting in a reduction in gaps by 61% in the 2DS portion (with 4,053 gaps remaining). In accordance with the Toronto agreement for pre-publication data sharing, comprehensive details of comparisons between our assembly and RefSeq v2.0 are not provided. Rather, we highlight and summarize a few noteworthy results from these comparisons.

Most notable are revisions to large structural misassemblies from RefSeq v1.0. Of 37 insertions, two were filled completely, while an additional 27 were partially filled or revised for improved accuracy in estimated gap size. As shown in the sample figures below, most of these updates are consistent with our assembly, thus providing independent evidence that our assembly workflow and resulting contigs are of high accuracy. Moreover, the T48 contig sequence (351 kb), which was unanchored in RefSeq v1.0, could be partially aligned to the 22.0-22.3 Mb region of the 2D chromosome in RefSeq v2.0. This result is consistent with the preliminarily identified location of the contig within the 20,8795-20,833 kb region of the 2D chromosome in RefSeq v1.0, determined using the physical map. In addition, all three inversions were completely corrected in RefSeq v2.0.

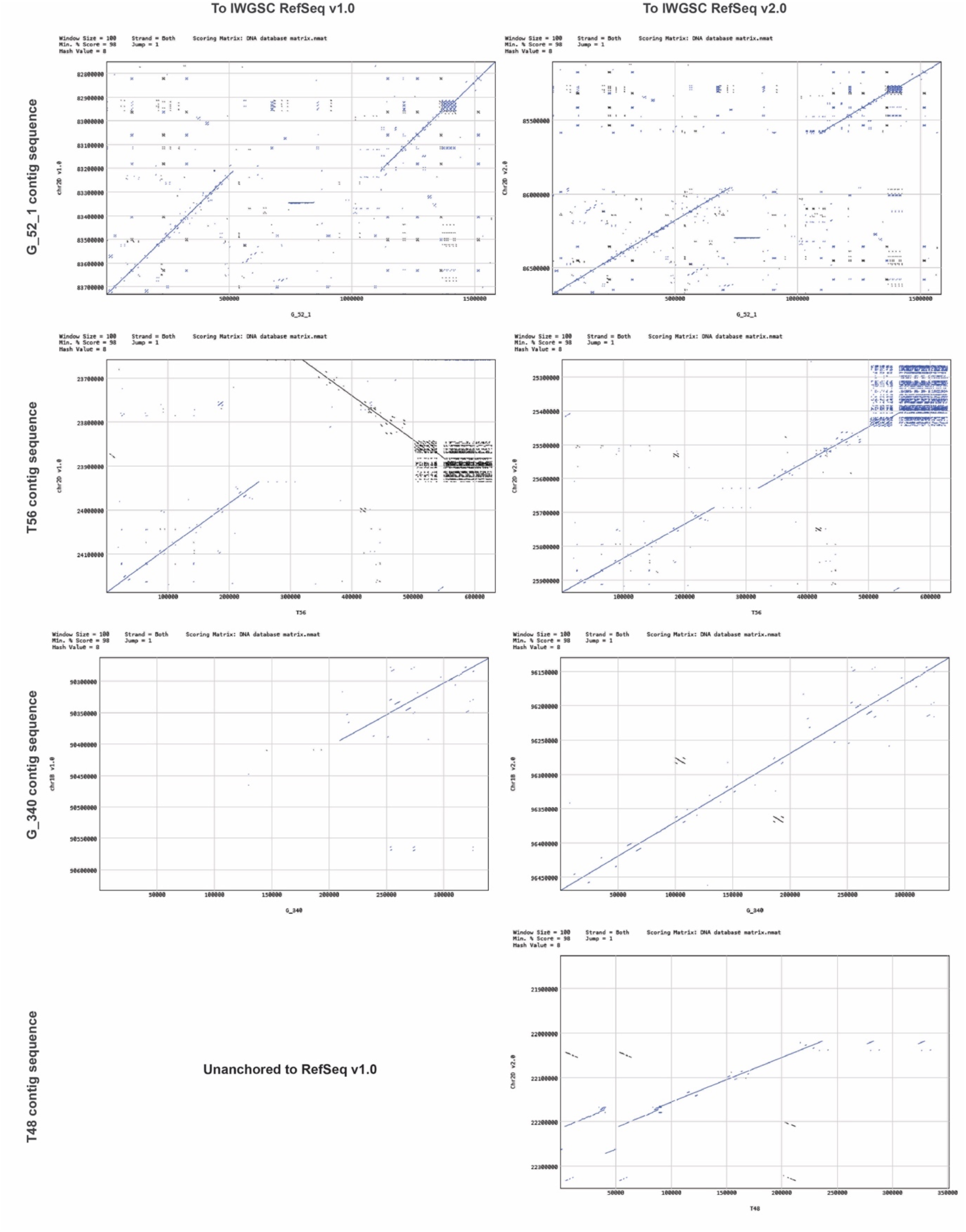

Our assembly features two large structural differences relative to RefSeq v2.0, which do not appear in comparison to RefSeq v1.0. These differences are located on chromosomes 2D and 1B. An inconsistency associated with a large segment (3.2 Mb) can be observed by dot-plot visualization of the G_142 contig sequence against the 2D chromosome reference sequences. This segment was relocated from the 268.0-271.2 Mb portion of Chr2D in RefSeq v1.0 to 260.4-263.7 Mb in RefSeq v2.0. As illustrated by the figures shown below, both our assembly and the pan-genome comparisons support the chromosome structure represented in RefSeq v1.0. Pan-genomic analysis was undertaken to characterize the collinearity of the G_142 contig sequence to corresponding portions of 10 wheat germplasms (http://www.10wheatgenomes.com), in addition to both RefSeq v1.0 and v2.0. These results indicate strong collinearity between G_142 and all samples except for RefSeq v2.0, for which relatively little collinearity was observed for the translocated portion.

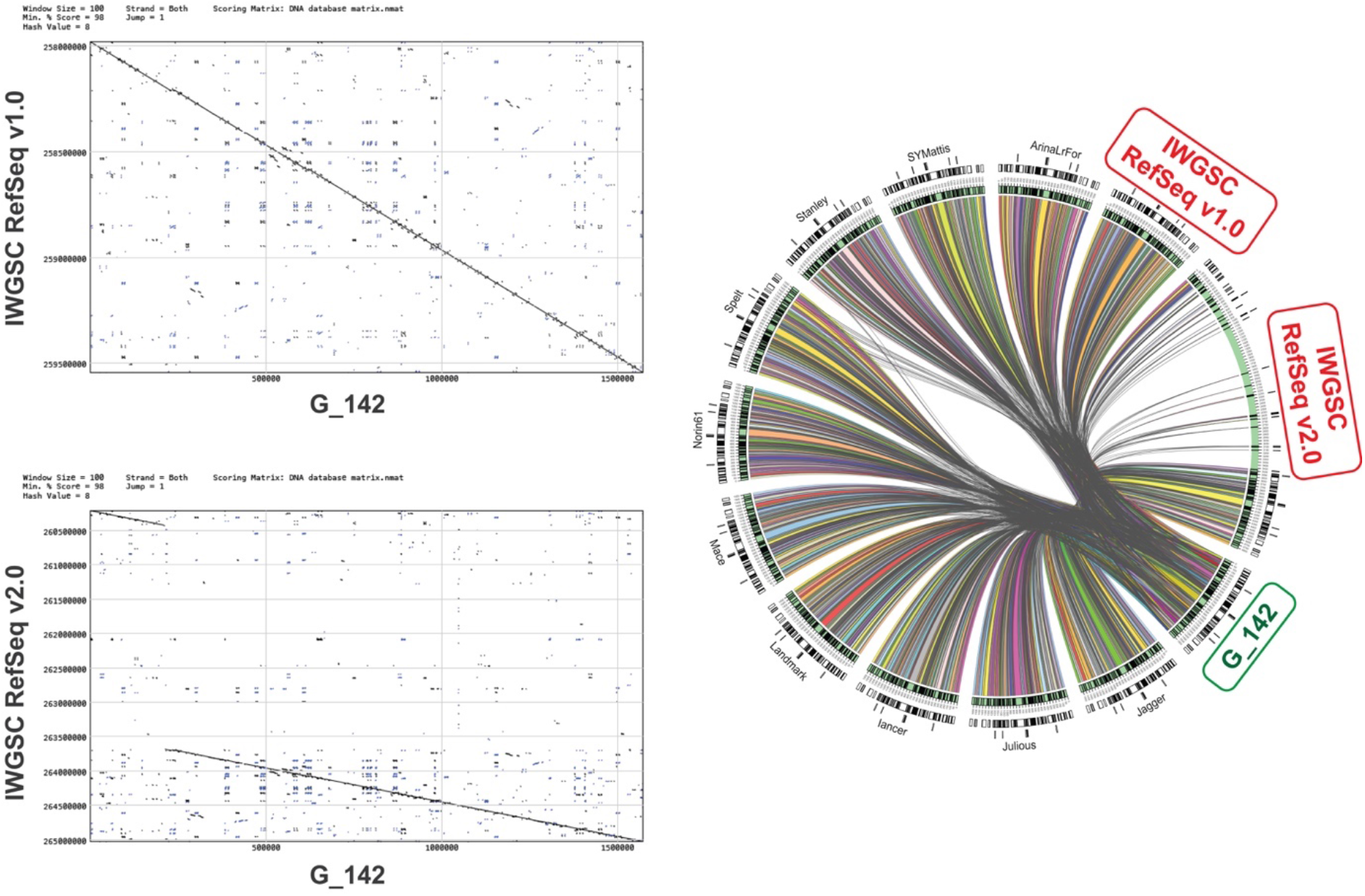

Upon integration of RefSeq v2.0 with our chromosome-scale contig sequences, a total of 3,632 gaps of the 2DS portion were filled completely, while 421 gaps remained unfilled.

**Supplemental Material 7.** Cost estimation and comparison to popular workflows.

Two examples are provided for comparison of estimated costs for our pooled hybrid sequencing design and Lamp assembler to other assembly workflows. The first example provides cost estimates for producing a reference sequence for a bread wheat germplasm other than Chinese Spring, while the second provides an estimate for simultaneous sequencing of multiple samples with relatively simple genome structures.

We first demonstrate the potential for application of our workflow towards producing a reference sequence for a bread wheat germplasm. To this end, two BAC libraries were constructed, each of which was composed of 250k clones with an average insert size of 150 kb. A total of 300k clones were sequenced to produce gapless BAC sequences covering the genome with 3× coverage. These BAC sequences were further assembled to chromosome-scale contigs with an expected average length exceeding 1.0 Mb. Contigs were anchored to an optical map, and their positions and relative distances were determined. Gaps were closed either by using consensus sequences of PacBio reads or by selective sequencing of clones from the un-sequenced portion of the BAC library.

To apply an assembly workflow similar to that used for the IWGSC RefSeq assembly, the initial chromosome-scale scaffolds were assembled from WGS short reads using either DeNovoMagic or TRITEX. Misassemblies were revised using WGP tags of BAC clones that were produced by a series of methods, including chromosome sorting, construction of chromosome-specific BAC libraries, and finally WGP tag sequencing. Several remaining misassemblies were further corrected by optical mapping. Gaps were filled by the filling step of our workflow, with a potential tenfold increase in costs resulting from the significantly increased number of gaps, as shown in the paper. The details of these comparisons are listed in the table below.

| Item | Estimated experimental costs (\$) for assembling the reference genome of a wheat germplasm by | |
| --- | --- | --- |
|  | our workflow | a workflow as used for IWGSC RefSeq |
| 2× 250k whole-genome BAC libraries (\$0.15 for each clone) | 75,000 | 0 |
| 300k BAC sequencing (\$5 for each clone) | 1,500,000 | 0 |
| 1,600 Gb whole-genome SMRT sequencing (20 Sequel II SMRT Cells, \$3000 for each) | 60,000 | 60,000 |
| Whole-genome Illumina sequencing (paired-ends and mate-pairs) | 0 | 30,000 |
| Sorting of all 21 chromosomes (\$50,000 for each) | 0 | 1,050,000 |
| 21 chromosome-specific BAC libraries (\$10,000 for each) | 0 | 210,000 |
| WGP for chromosome-specific libraries (\$150,000 for each) | 0 | 3,150,000 |
| Genome optical map | 100,000 | 100,000 |
| Correction of misassemblies and gaps | 150,000 | 1,500,000 |
| **Total** | **1,885,000** | **6,100,000** |

For the second example, we demonstrate the potential use of our workflow for assembling reference sequences for samples including 20 bacterial strains (each of ∼5 Mb in genome size), three monokaryon fungal strains (∼50 Mb each) and a rice germplasm (∼450 Mb). The Lamp assembler can be applied to sequence data produced by various workflows, as illustrated by this example. For each sample, a PE sequencing library with an average insert size of 350 bp was constructed, yielding short reads of approximately 1,000× genome coverage. A super pool was produced by mixing DNA solutions according to the desired coverage of long reads for each sample (80× for bacterial genomes, 130× for fungal and rice genomes). Long-read sequencing was performed using a single SMRT Cell on the Sequel II platform, which is expected to be sufficient for the super pool since a single SMRT Cell could be guaranteed by the service provider to have a minimum throughput of 80 Gb. For rice sequencing, contigs were joined to the chromosome-scale scaffold by optical mapping, and the few remaining gaps could be filled by assembly of at most 500 selected BAC clones.

Long-read-based non-hybrid assembly workflows have emerged as a common method in recently published studies. We estimated that up to four bacterial strains could be pooled together for high-coverage assembly using the sequencing results from a single SMRT Cell on the original Sequel platform. Each fungal strain was sequenced using an individual SMRT Cell on the original Sequel platform, while the rice germplasm was sequenced using one SMRT Cell on the Sequel II platform. Corrections of misassemblies can require dramatically increased costs relative to the initial assembly, as described in this paper. We estimate a threefold increase in cost associated with misassembly correction in our workflow. Details of our estimated cost comparisons across sequencing workflows are provided in the table below.

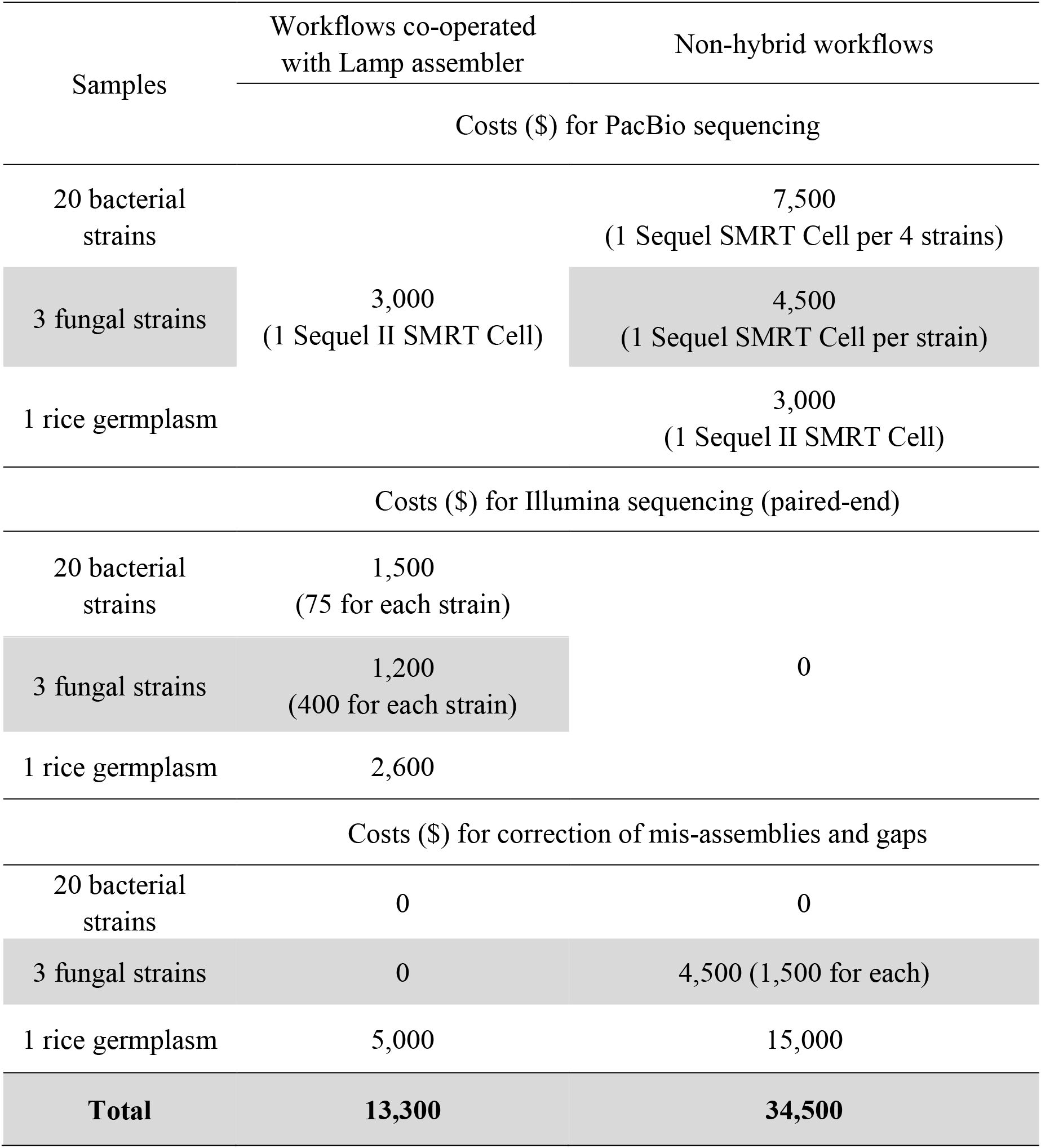

**Supplemental Material 8.** Details for construction of the 2DS-specific BAC library

## Plant material

Seeds of the double ditelosomic line of hexaploid *Triticum aestivum* L. cv. (‘Chinese Spring’ wheat) (2n = 20” + t”2DS + t”2DL) carrying the short and long arms of chromosome 2D in the form of telosomes were provided by Professor Adam J. Lukaszewski (University of California, Riverside, USA). The seeds were germinated in the dark at 25 ± 0.5 °C on moistened filter paper for 3 days to produce roots that were 2-3 cm in length. A total of 5,773 seeds were germinated in batches of 25-30 for the preparation of a total of 214 samples of suspensions of mitotic metaphase chromosomes.

## Preparation of chromosome suspensions

Mitotic chromosomes were isolated from synchronized root tip cells. Cycling root tip cells were first accumulated at the G1-S interphase by incubation of seedling root tips in 2 mM hydroxyurea at 25 ± 0.5 °C for 18 h. Samples were subsequently transferred to Hoagland’s solution and incubated for 5.5 h to recover from the hydroxyurea-mediated blockage of cell cycle progression. Next, mitotic cells were arrested in metaphase by incubation in 2.5 µM amiprophos-methyl for two h, followed by overnight ice-water treatment. Synchronized root tips were fixed in 2% (v/v) formaldehyde and Tris buffer solution at 5 °C for 20 min and then washed three times in Tris buffer, with each of the three washing steps performed for 5 min at 5 °C. Root tips were excised at 1 mm from the tip, and chromosomes were released by homogenization in 1 mL of LB01 nuclear lysis buffer using a Polytron PT1300D homogenizer (Kinematica AG, Littau, Switzerland) at 20,000 rpm for 13 s. Crude suspensions were filtered through a 50-µm pore nylon mesh to remove large tissue fragments.

## Flow cytometric analysis and sorting

Chromosome analysis and sorting were performed on a FACSVantage flow cytometer (Becton Dickinson, San José, USA) equipped with an argon-ion laser configured for multiline UV emission with an output power of 300 mW. A solution of 50 mM NaCl was used as the sheath fluid. Chromosome suspensions were stained with 4’,6-diamidino-2-phenylindole (DAPI) at a final concentration of 2 µg/mL, filtered through a 20-µm pore size nylon mesh and analysed at rates of 200-400 particles per second. DAPI fluorescence was measured using a fluorescence 1 (FL1) detector with a 424/44 bandpass filter. The relative fluorescence intensities of each chromosome suspension were recorded and plotted to produce histograms of the FL1 pulse area (FL1-A).

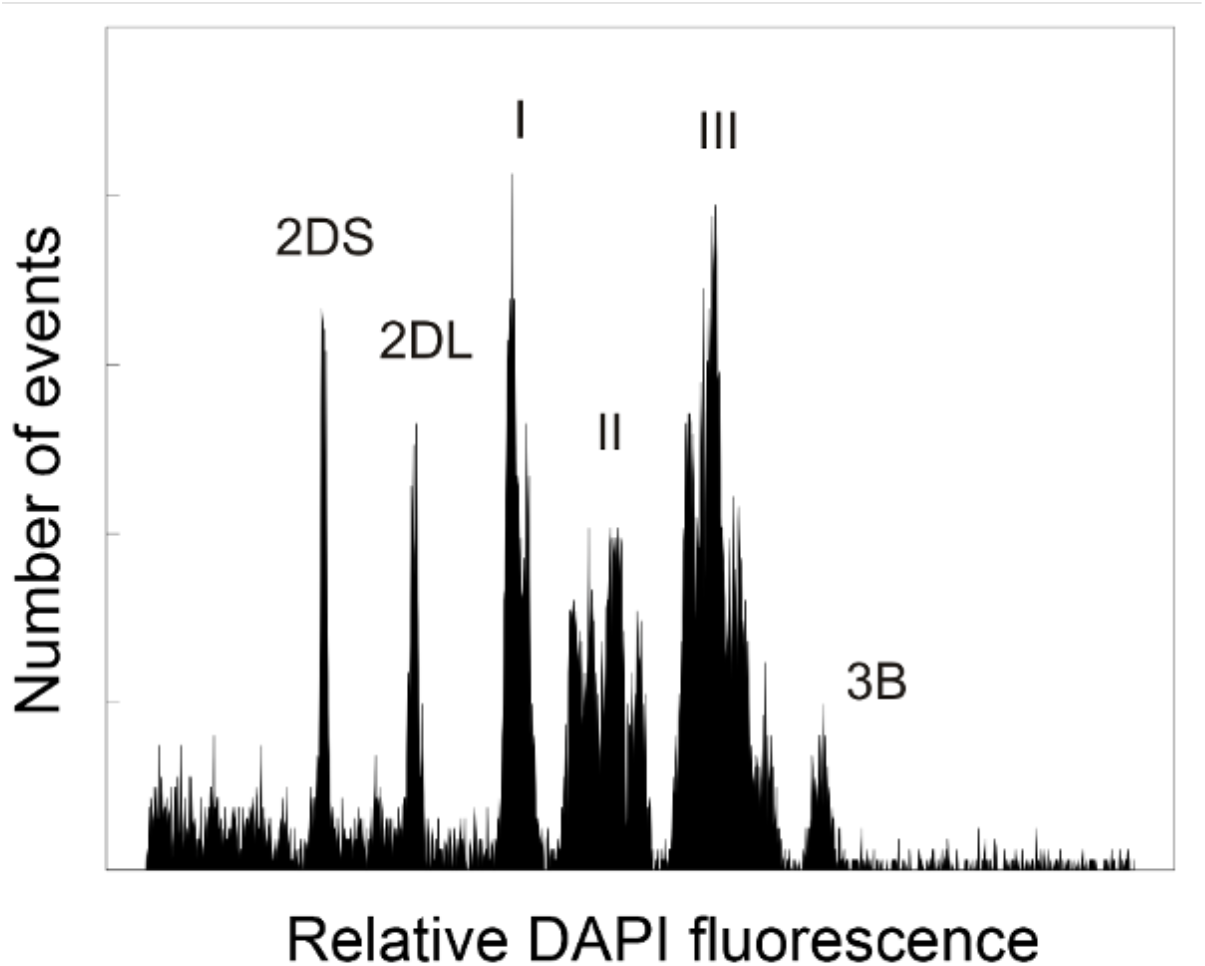

**Figure legend**. Histogram of relative fluorescence intensity (flow karyotype) obtained after the analysis of DAPI-stained mitotic metaphase chromosomes isolated from the double ditelosomic line of hexaploid wheat *Triticum aestivum* L. cv. Chinese Spring (2n = 20” + t”2DS + t”2DL), which carries the short and long arms of chromosome 2D in the form of telosomes 2DS and 2DL. The flow karyotype consists of a peak representing chromosome 3B, three clusters of peaks representing groups of chromosomes (I, II and III), and two peaks representing arms of chromosome 2D, which could be clearly distinguished, enabling isolation of the 2DS.

For chromosome sorting, gates were set on a dot-plot of FL1-A versus FL1 pulse width (FL1-W), and 2DS chromosomes were sorted at rates of 5-10 telosomes per second. A total of 7,750,000 2DS telosomes, corresponding to ∼5 μg of DNA, were flow-sorted in aliquots of ∼1.0 × 10^5^ into 160 μL of 1.5 × IB buffer. The identities and purities of the sorted chromosomes were determined microscopically after isolation via double FISH with probes for Afa and telomeric repeats. The average purity of sorted fractions was 88.26%, and the 2DS fractions were contaminated by a mix of other chromosomes, chromatids and chromosome arms.

## Preparation of high-molecular-weight (HMW) DNA

Flow-sorted chromosomes were pelleted at 200× g for 30 min at 4 °C and resuspended in 7.5 μL of 1× IB at 50 °C. Then, the samples were mixed with 4.5 μL of prewarmed 2% InCert low-melting-point agarose (GTG) in 1× IB. The mixture was poured into an 80-µL plug mould to form an agarose miniplug. The quality of HMW DNA was evaluated by pulsed-field gel electrophoresis (PFGE).

## Partial digestion, size selection and recovery of HMW DNA

Agarose miniplugs were washed twice for 1 h in 10:10 TE buffer (10 mM Tris, 10 mM EDTA). Subsequently, they were equilibrated on ice for 1 h in 10 mL of 1× *Hind*III buffer (Invitrogen) supplemented with 4 mM spermidine, 1 mM DTT and 0.1 mg/mL BSA. Partial *Hind*III digestion was performed for three 2DS miniplugs at a time, each using 6 conditions of *Hind*III enzyme concentration (0.02, 0.03, 0.05, 0.1, 0.2, and 1 units/tube) in 1 mL of buffer, incubated for 20 min at 37 °C. The samples were transferred to an ice bath, and digestion was terminated by the addition of 200 µL of 0.5 M EDTA stock solution at pH 8.0 and incubation of the resulting mixture for 30 min. Partially digested DNA was size-selected by PFGE with a 1% Gold SeaKem agarose (GTG) gel at 6 V/cm and 12 °C in 0.25× TBE for 14 h, with a 1.0 to 50 s switching interval and an angle of 120°. After electrophoresis, the edges of the gel containing size markers were excised and stained with ethidium bromide.

Five regions of the gel (100-150, 150-200 and 200-250 kb) were excised and equilibrated with 1.3× TAE buffer twice for 1 h each. The size-selected DNA was isolated by electroelution in 1.3× TAE using a BioRad Electroelution system (Model 422). The optimum electroelution (electrophoresis) time for concentrating the partially digested DNA in the lower-39 µL fraction, directly on the membrane CAPS, was determined by conducting several electrophoresis monitoring experiments with whole genomic DNA eluted over a range of run times. The DNA concentration was estimated in 1% normal agarose gels using 5 µL of electroeluted DNA and a dilution series of *Hind*III-digested lambda DNA as a standard.

## Ligation and transformation

The complete remaining DNA fraction (approx. 33 µL) was ligated at 16 °C overnight in a 50-µL reaction with 4 units of T4 DNA ligase (Invitrogen) and 4 ng of pIndigoBAC *Hind*III Cloning Ready vector (Epicentre) prepared for high-efficiency cloning by digestion with *Hind*III. The ligation mix was incubated at 4 °C for 90 min. Fifteen microlitres of the resulting ligation was added to 110 µL of MegaX DH10B T1 electrocompetent cells, which were then incubated at room temperature for 5 min. The incubated mixture was divided into 15-µL aliquots, each of which was electroporated using a Gibco BRL Cell-Porator System (Life Technologies) with the following settings: 350 V, 330 μF capacitance, low ohm impedance, fast charge rate, and 4 kΩ resistance. The electroporation solutions were pooled in a tube containing 3 mL of recovery media and incubated for 60 min at 37 °C and 175 rpm on an orbital shaker. Aliquots of the recovery media with recombinant cells were plated on LB plates containing 12.5 μg/mL chloramphenicol, 50 μg/mL X-Gal and 25 μg/mL IPTG. The plates were incubated at 37 °C overnight. A Q-bot (Genetix) was used to identify recombinant colonies by their white phenotype, and these colonies were picked and used to inoculate wells of 384-well plates containing 90 µL of LB freezing buffer (Peterson *et al*. 2000). The plates were incubated overnight at 37 °C, triplicated and stored at –80 °C. The complete 2DS-specific BAC library (code TaaCsp2DShA) comprises 43,008 clones ordered in 112 384-well plates.

## Isolation of BAC DNA and insert analysis

Individual BAC clones were cultured overnight in deep 96-well plates with wells containing 1.5 mL of LB supplemented with 12.5 µg/mL chloramphenicol. BAC DNAs were isolated and digested to completion with *Not*I. DNA fragments were size-separated by PFGE in a 1% Gold SeaKem agarose (GTG) gel at 6 V/cm, with a 1-15 s switch time ramp and an angle of 120°, for 14 h at 14.0 °C in 0.25× TBE buffer. Analysis of 120 BAC clones indicated that the 2DS BAC library had an average insert size of 132 kb. Almost two-thirds of the 2DS library (62.5%, in plate Nos. 1-70) was constructed from a fraction with an average insert size of 126 kb. The second fraction of clones used for the 2DS library construction (37.5%, in plate Nos. 71-112) had a greater average insert size of 142 kb.

## Coverage and specificity of the BAC library

The coverage for 2DS was estimated as 15.6× based on the average insert size, the total number of clones and an 11.74% rate of contamination by other chromosomes. The total size of 2DS was calculated to be 316 Mb based on the relative length of 2DS (1.86%) compared to the nuclear genome size of common wheat (16,974 Mb/1C). Accounting for the size of 2DS and the 11.74% rate of contamination by undesired chromosomes in the 2DS library, the probability of a given DNA sequence in the library originating from 2DS was determined to be 99.6%.

